# Shedding Light on the D_1_-Like Receptors: A Fluorescence-Based Toolbox for Visualization of the D_1_ and D_5_ Receptors

**DOI:** 10.1101/2023.09.25.559386

**Authors:** Niklas Rosier, Denise Mönnich, Martin Nagl, Hannes Schihada, Alexei Sirbu, Nergis Konar, Irene Reyes-Resina, Gemma Navarro, Rafael Franco, Peter Kolb, Paolo Annibale, Steffen Pockes

## Abstract

Dopamine D_1_-like receptors are the most abundant type of dopamine receptors in the central nervous system and, even after decades of discovery, still highly interesting for the study of neurological diseases. We herein describe the synthesis of a new set of fluorescent ligands, structurally derived from D_1_R antagonist SCH-23390 and labeled with two different fluorescent dyes, as tool compounds for the visualization of D_1_-like receptors. Pharmacological characterization in radioligand binding studies identified UR-NR435 (**25**) as a high-affinity ligand for D_1_-like receptors (p*K_i_* (D_1_R) = 8.34, p*K_i_* (D_5_R) = 7.62) with excellent selectivity towards D_2_-like receptors. Compound **25** proved to be a neutral antagonist at the D_1_R and D_5_R in a G_s_ heterotrimer dissociation assay, an important feature to avoid receptor internalization and degradation when working with whole cells. The neutral antagonist **25** displayed rapid association and complete dissociation to the D_1_R in kinetic binding studies using confocal microscopy verifying its applicability for fluorescence microscopy. Moreover, molecular brightness studies determined a single-digit nanomolar binding affinity of the ligand, which was in good agreement with radioligand binding data. For this reason, this fluorescent ligand is a useful tool for a sophisticated characterization of native D_1_ receptors in a variety of experimental setups.

## Introduction

Dopamine (DA, Figure 1) is a catecholaminergic neurotransmitter which is widely expressed in the central nervous system (CNS), especially in the nigrostriatal, mesolimbic, mesocortical and tuberoinfundibular systems.[1–3] DA exerts its physiological functions via five G protein-coupled receptors (GPCRs), the dopamine D receptors (D_1-5_R).[4] These five dopamine receptors are divided into two sub-families based on their ability to activate or inhibit the adenylyl cyclase and hence the production of the second messenger cAMP.[5] The D_1_R and D_5_R are Gα_s_-coupled and form the D_1_-like receptor family, whereas the D_2_R, D_3_R and D_4_R are Gα_i/o_-coupled and members of the D_2_-like receptor family.[6,7] Due to the wide expression of dopamine and its receptors in the central and peripheral nervous system they have been in the focus of research for many years. Malfunctions of the dopaminergic systems are related to a large number of diseases including schizophrenia, Parkinson’s disease (PD), bipolar disorder, depression, restless leg syndrome, hyperprolactinaemia, hypertension, gastroparesis, nausea, and erectile dysfunction.[4,5,8–13] Most of the drugs that are on the market target the D_2_-like receptors, such as haloperidol, pramipexol and metoclopramide.[14–16] The only approved drug targeting the D_1_-like receptors is the peripheral acting partial agonist fenoldopam for the intravenous treatment of hypertension.[17,18] The central active D_1_R selective full agonist dihydrexidine (Figure 1) was tested in clinical trials for the treatment of PD, but failed due to severe side effects.[19,20] Despite decades of research, the complex dopaminergic system including the interactions of dopamine receptors with others, such as other dopamine receptors, histamine, NMDA or adenosine receptors is not fully understood yet and further investigations are needed.[21–24]

**Figure 1.**
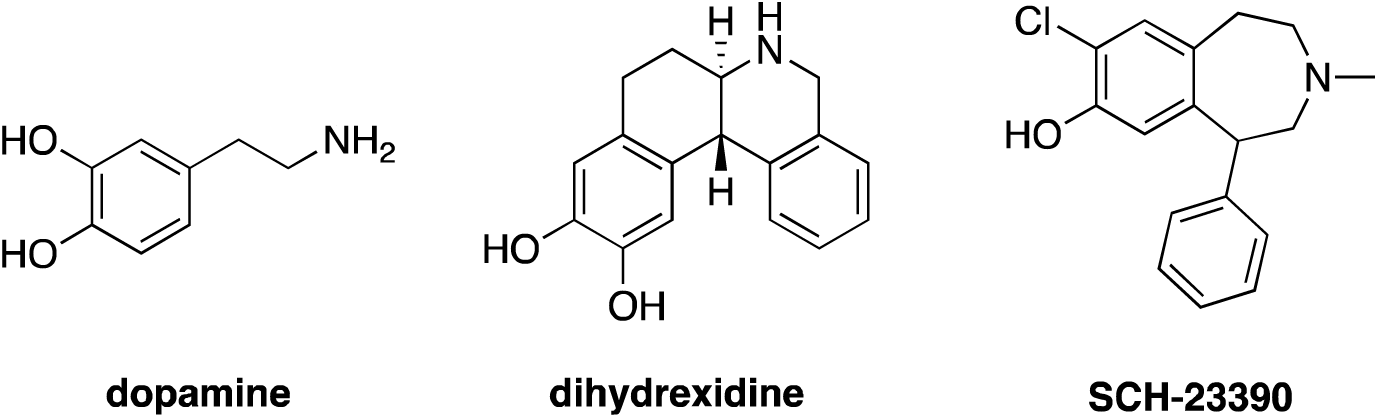
Chemical structures of dopamine, dihydrexidine (D_1_R agonist) and SCH-23390 (D_1_R/D_5_R antagonist).

Fluorescent ligands represent attractive alternatives to radioligands, for example, to visualize receptors in cells and tissues or for studying the effect of drug-receptor-interactions with advanced biophysical technologies.[25,26] Many fluorescence-based techniques have been developed to study ligand-protein-interactions, including flow cytometry, fluorescence anisotropy, fluorescence polarization as well as resonance energy transfer (RET) based assays.[27–29] Furthermore fluorescence microscopy techniques such as confocal microscopy and total internal reflection fluorescence (TIRF) microscopy, are often used for these purposes.[30,31]

We herein report the development and validation of a series of six novel fluorescent ligands for D_1_-like receptors and their use for binding studies in fluorescence microscopy. Although several fluorescent ligands for the D_1_R have previously been published, to our knowledge none of them was tested at the D_5_R or has been used for molecular brightness studies.[32–34] The goal of the study is to find useful pharmacological tools that can further help to explore the complex functionalities and interactions of D_1_-like receptors.

## Results and Discussion

### Design rationale

Fluorescent ligands can be considered as an entity of three distinct parts: the pharmacophore, the linker and the fluorescent dye. Each of the three parts must be chosen individually based on the intended use of the tool compound.[35–37]

The known D_1_R/D_5_R antagonist SCH-23390 (Figure 1) was chosen as the structural motif for the pharmacophore because it displays high affinity at the D_1_R and D_5_R and a high selectivity within the dopamine receptor family.[38] As a pharmacophore, an antagonistic partial structure was preferred since agonists can induce receptor internalization and degradation, which could be disadvantageous for binding studies.[35] Structure-activity relationships (SAR) revealed that structural modifications and attachments at the *para*-position of the phenyl group are well tolerated and that a sulfonamide group may be beneficial for binding affinity.[39,40] In accordance with these SAR results, cryo-EM structures of D_1_R-Gs complexes bound to an agonistic SCH-23390-derivative, SKF-83959, showed that this position points towards the extracellular space and is not involved in ligand-receptor interaction.[41] Therefore, this position and chemical moiety were chosen for the attachment of the linker.

Based on previously described D_1_R fluorescent ligands, which had the fluorescent dye directly attached to the pharmacophore, two ligands with a short alkylic linker were synthesized.[34] Additionally, fluorescent ligands with longer PEG-based linkers were prepared which should allow the fluorophore to reach outside the binding pocket and therefore, should have reduced impact on the ligand binding. PEG-based linkers are commonly used in fluorescent ligands because they usually don’t interact with cell membranes, are chemically stable and well soluble in water.[42] The connection of the pharmacophore with the PEG-unit was performed in three steps. First a short alkylic spacer was attached to the pharmacophore which was then connected via a peptide coupling reaction to propargylamine. This product was coupled via a copper catalyzed azide alkyne click (CuAAC) reaction to a PEG-2 or PEG-3 unit, which was orthogonally functionalized with a primary amine and an azide group (**17a**, **17b**). The additional triazole moiety increases the solubility of the final fluorescent ligands.

The choice of the fluorescent dye is crucial and depends mainly on the intended use of the final fluorescent ligand. Important criteria to consider are spectral properties, quantum yield, solubility, solvatochromic effects, chemical stability, photostability, commercial availability, and an efficient coupling to the pharmacophore.[35] The 5-TAMRA fluorescent dye was chosen to label the ligands because it is known to yield excellent results in fluorescence microscopy.[43–45] Additionally, DY549-P1 from Dyomics was used, as it possesses similar spectral properties as 5-TAMRA. Both dyes are hydrophilic, reducing the risk of unspecific interactions with cell membranes as compared to, for example, BODIPY fluorescent dyes, leading to lower non-specific binding.[46]

### Chemistry

The syntheses of the pharmacophore and linkers were performed separately followed by the connection of both parts. The last step in the synthesis of the different fluorescent ligands was always the labeling with a fluorescent dye.

The pharmacophore was synthesized following the general synthetic route for benzazepines used by Neumeyer et al. and Shen et al. with modifications (Scheme 1).[47,48] In a three-step synthesis 4-nitroacetophenone was transformed into the corresponding epoxide **3** with good yields. Commercially available 4-methoxyaldehyde was first chlorinated using sulfurylchloride in acetic acid to afford 3-chloro-4-methoxybenzaldehyde (**4**). A Henry-reaction with nitromethane in acetic acid, followed by a reduction with LiAlH_4_ in THF afforded the primary amine **6**. The secondary amine **8** was prepared via the introduction of a Boc-protecting group (**7**) and subsequent reduction with LiAlH_4_. This two-step synthesis provided selectively the mono-methylated amine in high yields. A nucleophilic substitution of the secondary amine **8** with epoxide **3** in acetonitrile generated product **9** in excellent yields which was converted to the benzazepine **10** by an acid catalyzed cyclization reaction using Eaton’s reagent at room temperature for 72h. After deprotection of the methyl-group with boron tribromide in DCM, the resulting phenol **11** was protected with acetyl chloride or triisopropylsilyl chloride, respectively (**12a**, **12b**). A reduction of the nitro group with hydrogen and catalytical amount of palladium on charcoal in a mixture of MeOH and THF afforded the anilines **13a** and **13b**.

**Scheme 1.**
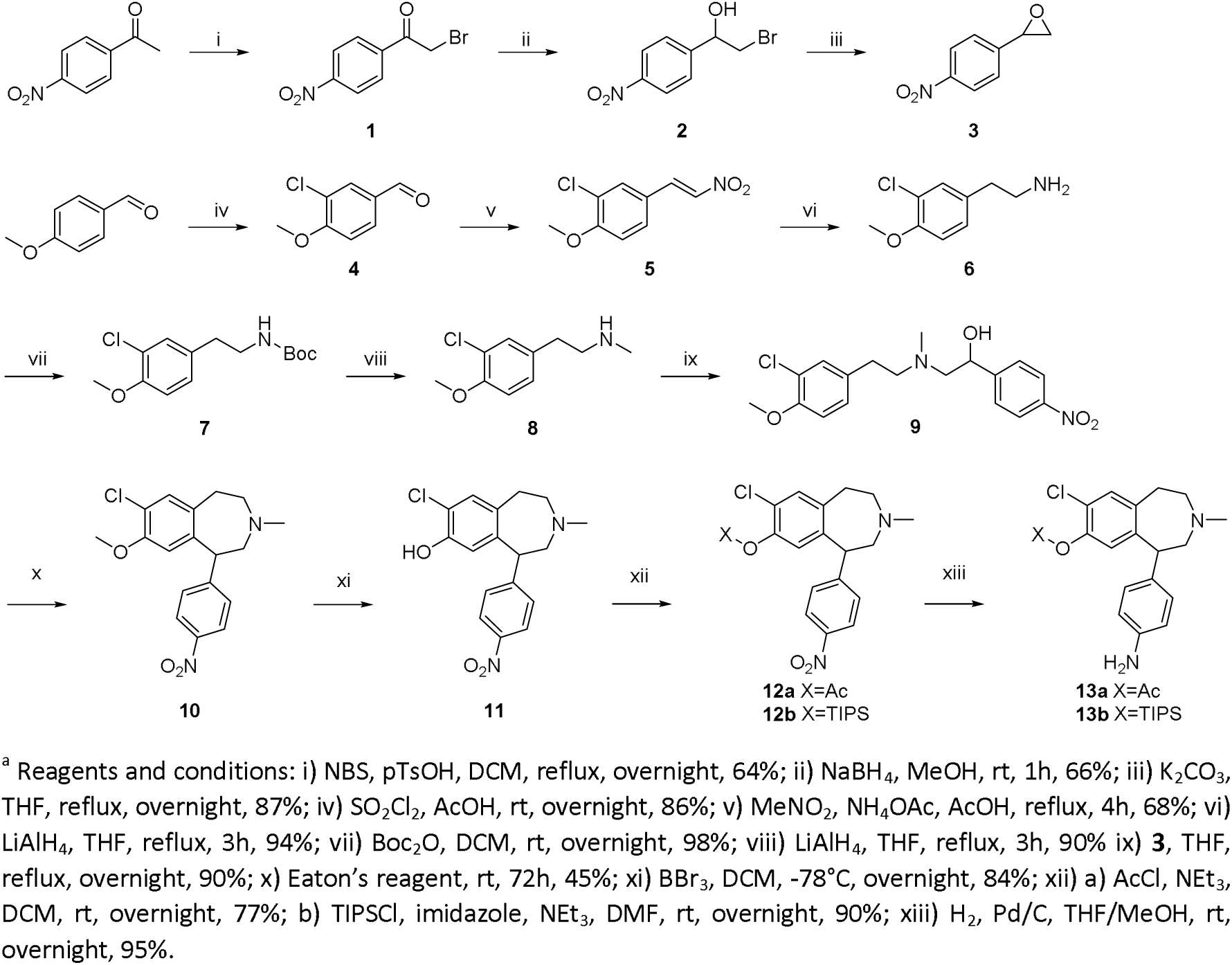
Synthesis of the pharmacophores **13a** and **13b**

For the introduction of the sulfonylamide moiety two different sulfonyl chlorides (**14a**, **14b**) were required. They were synthesized starting from aliphatic bromides using thiourea and *N*-chlorsuccinimide (Scheme 2A) following the procedure of Yang and Xu et al.[49] The PEG-linkers (**22a**, **22b**) were synthesized starting from the corresponding polyethylene glycols following the reaction protocol published by Iyer et al.[43,50]

**Scheme 2.**
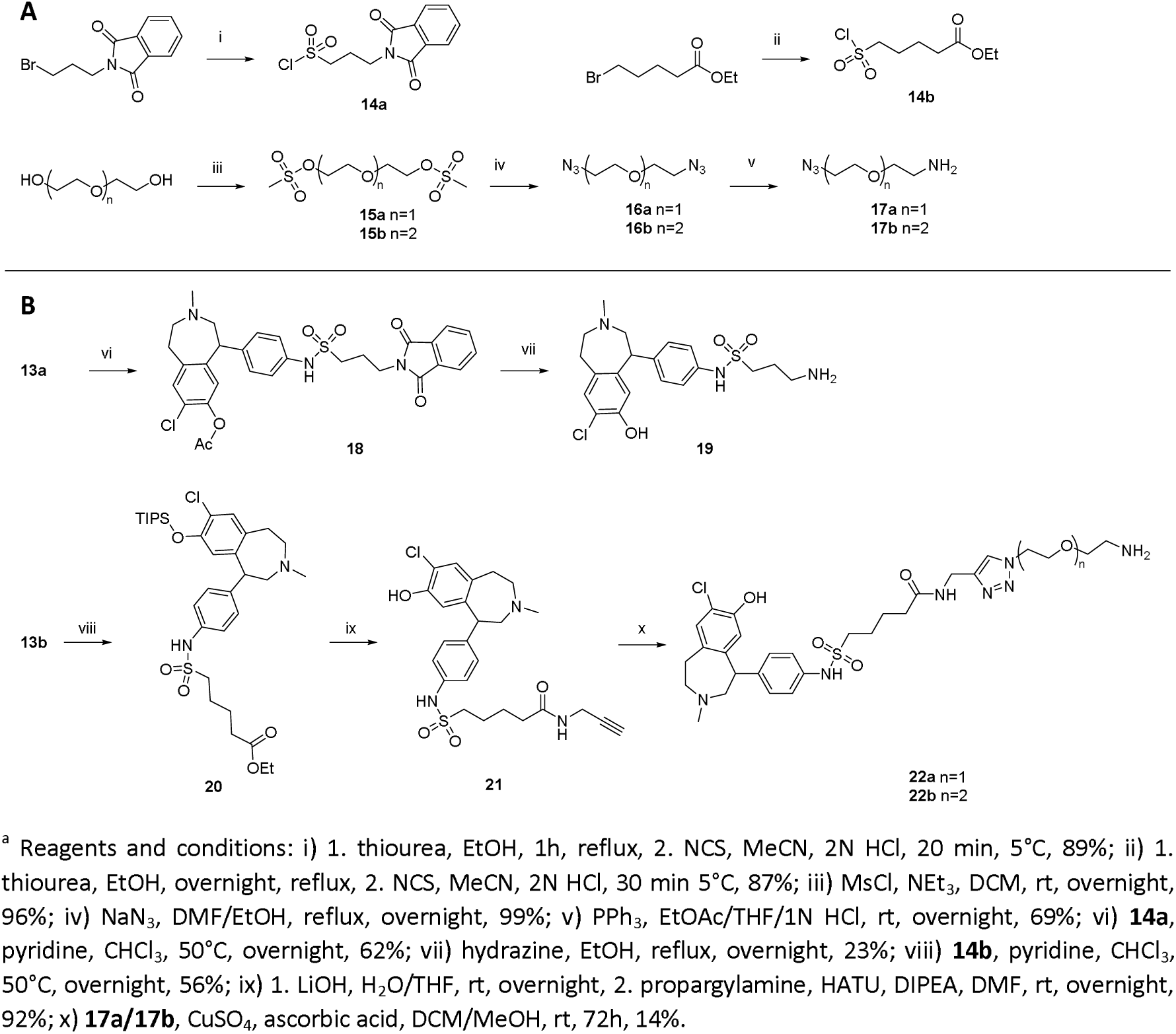
Synthesis of the sulfonylchlorides and linkers (**A**) and preparation of the final precursors (**B**)^a^

Coupling of the sulfonylchlorides **14a** and **14b** with the anilines **13a** and **13b** in the presence of pyridine afforded the structures **18** and **20** (Scheme 2B) in moderate yields. **18** was converted to the primary amine **19** using hydrazine hydrate in ethanol which simultaneously cleaved the acetyl group affording the free phenol. Purification by preparative HPLC afforded **19** in high purity as the final precursor for coupling to the fluorescent dyes. Ester hydrolysis of **20** in basic conditions and subsequent amide coupling with propargylamide using HATU as a coupling reagent obtained the terminal alkyne **21** in high yields. The triisopropylsilyl protecting group was cleaved as well during this reaction affording the free phenol which is essential for ligand binding to the D_1_-like receptors. By using a copper catalyzed alkyne azide click reaction protocol with CuSO_4_ pentahydrate and ascorbic acid as catalysts **21** was coupled to the PEG linkers. Due to complex purification by preparative HPLC the products (**22a, 22b**) were isolated in low but sufficient yields for the coupling with the fluorescent dyes.

All three precursors were labelled with both fluorescent dyes (5-TAMRA-NHS ester and DY549-P1-NHS ester) in DMF with an excess of NEt_3_ as a base (Scheme 3). Purification by preparative HPLC afforded the final fluorescent ligands (**23**-**28**) in moderate to good yields, great purity and stability (Figure S1-S6; Supporting Information (SI)).

**Scheme 3.**
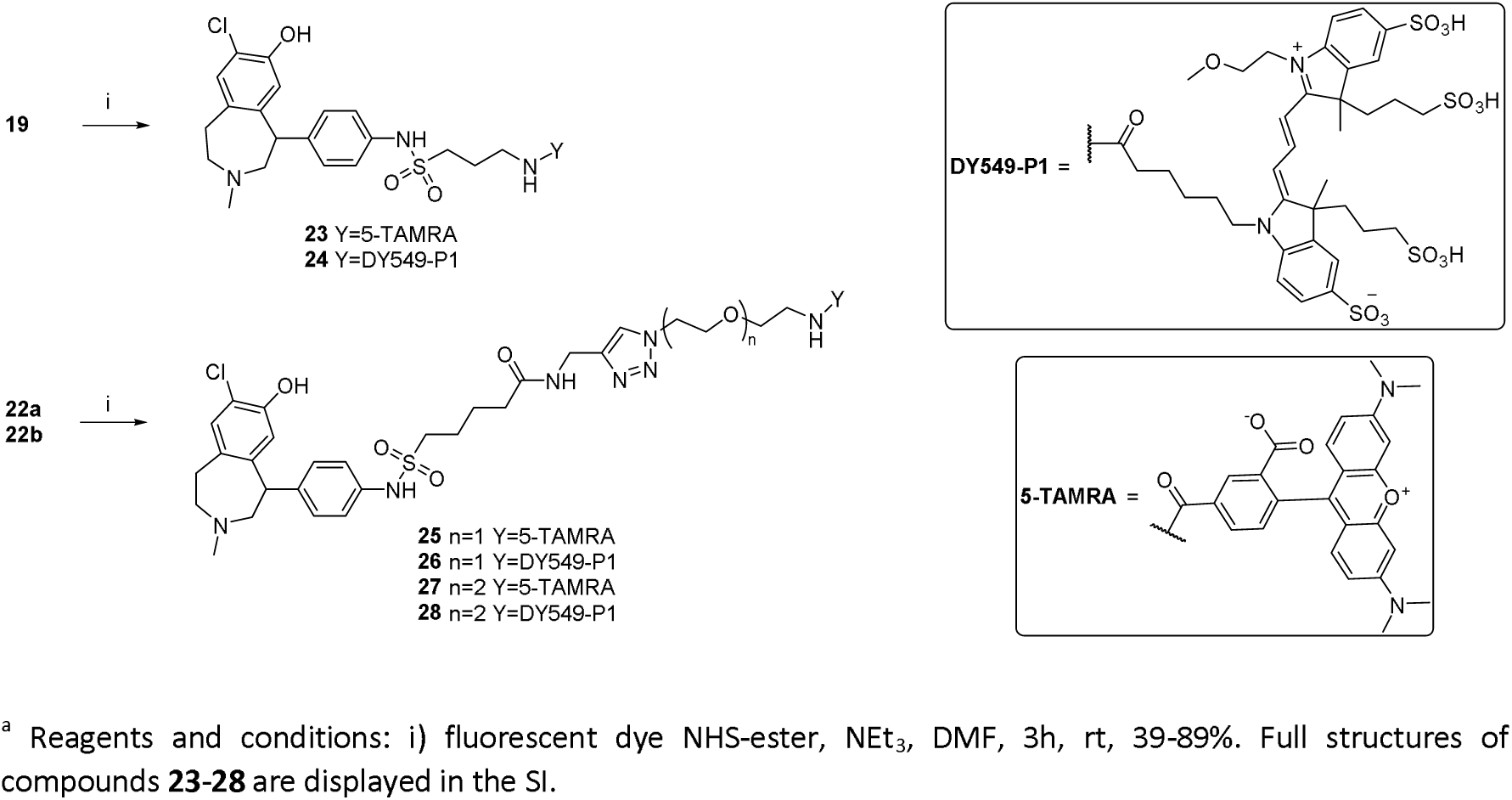
Synthesis of the fluorescent ligands **23-28^a^**

### Pharmacological Characterization

To pharmacologically characterize the new set of fluorescent ligands, we first determined their D_1_-like receptor affinity and picked the best compounds to test for selectivity within the dopamine receptor family. Therefore, all compounds were initially tested in radioligand competition binding assays at the wild type D_1_R (Figure 2A). Here, the fluorescent ligands **24**, **25** and **27** showed the highest affinities, with the two TAMRA derivatives **25** (p*K_i_* (D_1_R) = 8.34, Table 1) and **27** (p*K_i_* (D_1_R) = 8.02, Table 1) showing single-digit nanomolar affinities. Since the PEG-containing compounds **25-28** were able to show significantly better properties in terms of solubility, further competition experiments were performed with the same compounds at the D_5_R (Figure 2B). In all cases, reduced affinity values of about 0.4-0.8 logarithmic units were found. Again, the best results were obtained with **25** (p*K_i_* (D_5_R) = 7.62, Table 1) and **27** (p*K_i_* (D_5_R) = 7.65, Table 1). Both compounds were subsequently tested at the D_2long_R, D_3_R, and D_4_R, and only weak binding to the D_2_-like receptors was detected, representing a great selectivity profile toward D_2_-like receptors for **25** and **27** (at least 1,000-fold for D_1_R and at least 500-fold for D_5_R). Because of the highest affinity and selectivity, **25** seemed to be the most promising ligand to be used as a fluorescent tracer in microscopy studies. For this purpose, knowing the ligand’s mode of action is of great importance since agonists can alter the results of binding studies by inducing receptor internalization and degradation. Therefore, we confirmed the antagonistic behavior of **25** in a BRET-based G_s_ heterotrimer dissociation assay (G_s_-CASE) (Figure 3; Table 2).[51] In the agonist mode, only at high concentrations of 1 μM a slight decrease could be observed, which is most likely due to optical interference of **25** with the BRET components of G_s_-CASE as observed in an earlier study with a 5-TAMRA-labeled histamine H_3_ receptor ligand (Figure 3A).[43] In contrast, **25** reduced already at lower concentration the 1 μM dopamine-induced BRET response (p*K_b_* (D_1_R) = 7.29, p*K_b_* (D_5_R) = 9.48; Table 2), indicating that it acts as a neutral antagonist at both, D_1_ and D_5_ receptors. Concentration-response curves of dopamine-induced G_s_ activation at the D_1_R and D_5_R are shown in Figure S7 (SI).

**Figure 2.**
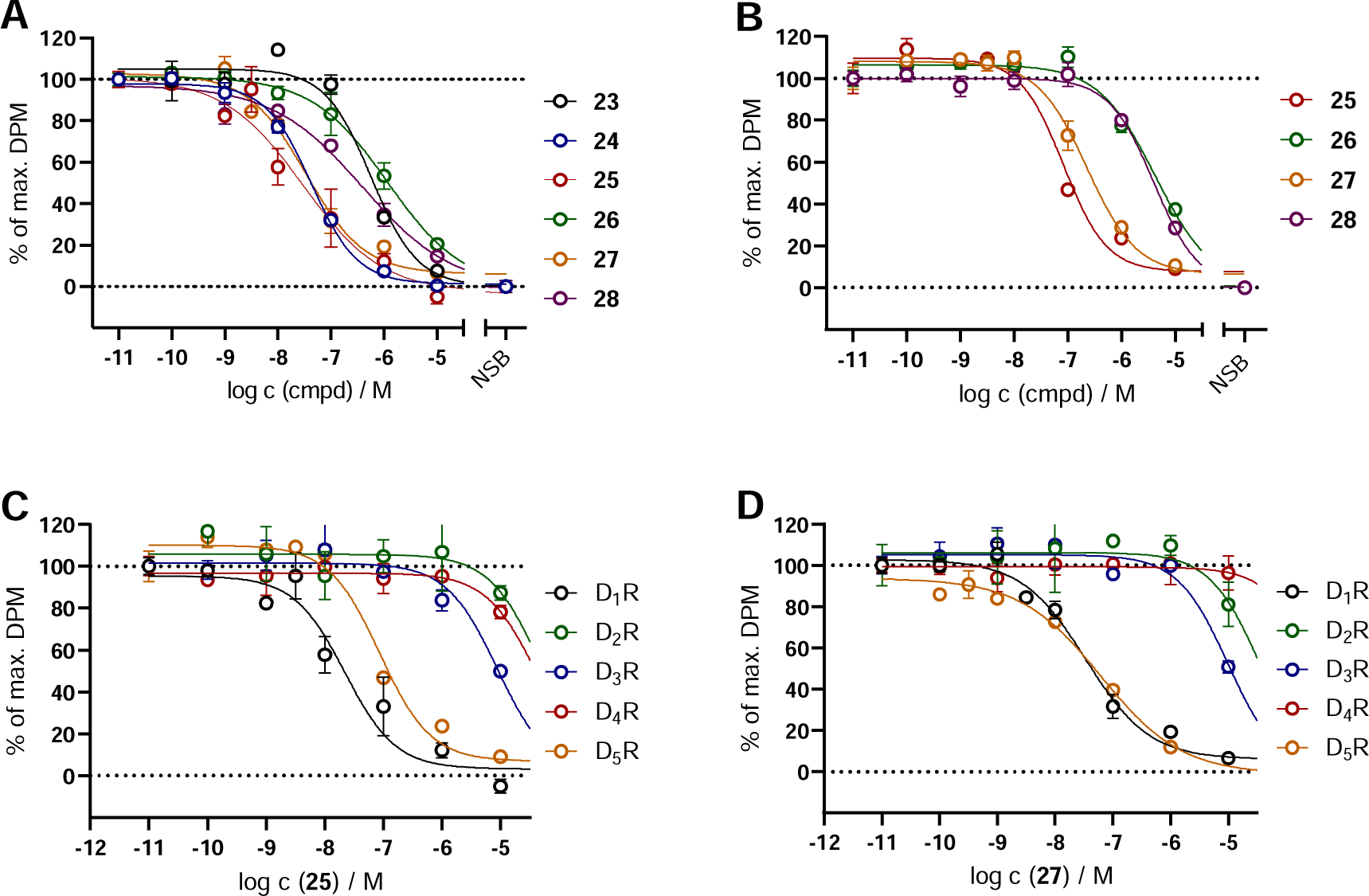
Displacement curves from radioligand competition binding experiments at the D_1_R (**A**) and D_5_R (**B**) homogenates performed with compounds **23**-**28** and the respective radioligands (cf. Table 1 footnotes). Displacement curves of **25** (**C**) and **27** (**D**) at the D_1-5_R homogenates. Graphs represent the means from three independent experiments each performed in triplicates. Data were analyzed by nonlinear regression and were best fitted to sigmoidal concentration-response curves.

**Table 1.** Binding data of **23-28** at the dopamine receptors.

| compound | Radioligand competition binding<sup>a</sup> |  |  |  |  |  |  |  |  |  |
| --- | --- | --- | --- | --- | --- | --- | --- | --- | --- | --- |
|  | pK<sub>i</sub> |  |  |  |  |  |  |  |  |  |
|  | D<sub>1</sub>R<sup>b</sup> | N | D<sub>2long</sub>R<sup>c</sup> | N | D<sub>3</sub>R<sup>d</sup> | N | D<sub>4</sub>R<sup>e</sup> | N | D<sub>5</sub>R<sup>f</sup> | N |
| **23** | 6.40 ± 0.39 | 3 | n.d. | - | n.d. | - | n.d. | - | n.d. | - |
| **24** | 7.99 ± 0.18 | 3 | n.d. | - | n.d. | - | n.d. | - | n.d. | - |
| **25** | 8.34 ± 0.21 | 3 | < 5 | 3 | 5.67 ± 0.06 | 3 | < 5 | 3 | 7.62 ± 0.03 | 3 |
| **26** | 6.54 ± 0.23 | 3 | n.d. | - | n.d. | - | n.d. | - | 6.03 ± 0.12 | 3 |
| **27** | 8.02 ± 0.14 | 3 | < 5 | 3 | 5.57 ± 0.03 | 3 | < 5 | 3 | 7.65 ± 0.14 | 3 |
| **28** | 6.85 ± 0.07 | 3 | n.d. | - | n.d. | - | n.d. | - | 6.06 ± 0.10 | 3 |

<sup>a</sup>Competition binding assay at HEK293T\_CRE Luc 2P D<sub>1</sub>R homogenates, HEK293T\_CRE Luc 2P D<sub>2long</sub>R homogenates, HEK293T\_CRE Luc 2P D<sub>3</sub>R homogenates, HEK293T\_CRE Luc 2P D<sub>4</sub>R homogenates, or HEK293T\_CRE Luc 2P D<sub>5</sub>R homogenates. <sup>b</sup>Displacement of 1 nM [<sup>3</sup>H]SCH-23390 (*K<sub>d</sub>* = 0.3 nM). <sup>c</sup>Displacement of 0.05 nM [<sup>3</sup>H]*N*-Methylspiperone (*K<sub>d</sub>* = 0.015 nM). <sup>d</sup>Displacement of 0.05 nM [<sup>3</sup>H]*N*-Methylspiperone (*K<sub>d</sub>* = 0.026 nM). <sup>e</sup>Displacement of 0.1 nM [<sup>3</sup>H]*N*-Methylspiperone (*K<sub>d</sub>* = 0.078 nM). <sup>f</sup>Displacement of 1 nM [<sup>3</sup>H]SCH-23390 (*K<sub>d</sub>* = 0.4 nM). Data shown are mean values ± SEM of N experiments, each performed in triplicates. Data were analyzed by nonlinear regression and were best fitted to sigmoidal concentration-response curves. Displacement curves are shown in Figure 2.

**Table 2.** Functional data of **25** at the D_1_R and D_5_R.

| compound | Gα<sub>s</sub> activation<sup>a</sup> |  |  |  |
| --- | --- | --- | --- | --- |
|  | pK<sub>b</sub> |  |  |  |
|  | D<sub>1</sub>R<sup>b</sup> | N | D<sub>5</sub>R<sup>c</sup> | N |
| **25** | 7.52 ± 0.09 | 3 | 9.60 ± 0.16 | 3 |

<sup>a</sup>Competition binding experiment at HEK293 cells expressing the G<sub>s</sub> BRET sensor with the wild type D<sub>1</sub>R or D<sub>5</sub>R. <sup>b</sup>Inhibition of dopamine-induced (c = 1 μM, EC<sub>50</sub> = 393 nM, Figure S7, SI) G<sub>s</sub> activation. <sup>c</sup>Inhibition of dopamine-induced (c = 1 μM, EC<sub>50</sub> = 26 nM, Figure S7, SI) G<sub>s</sub> activation. Data shown are mean values ± SEM of N experiments, each performed in duplicates. Data were analyzed by nonlinear regression and were best fitted to sigmoidal concentration-response curves. Displacement curves are shown in Figure 3.

**Figure 3.**
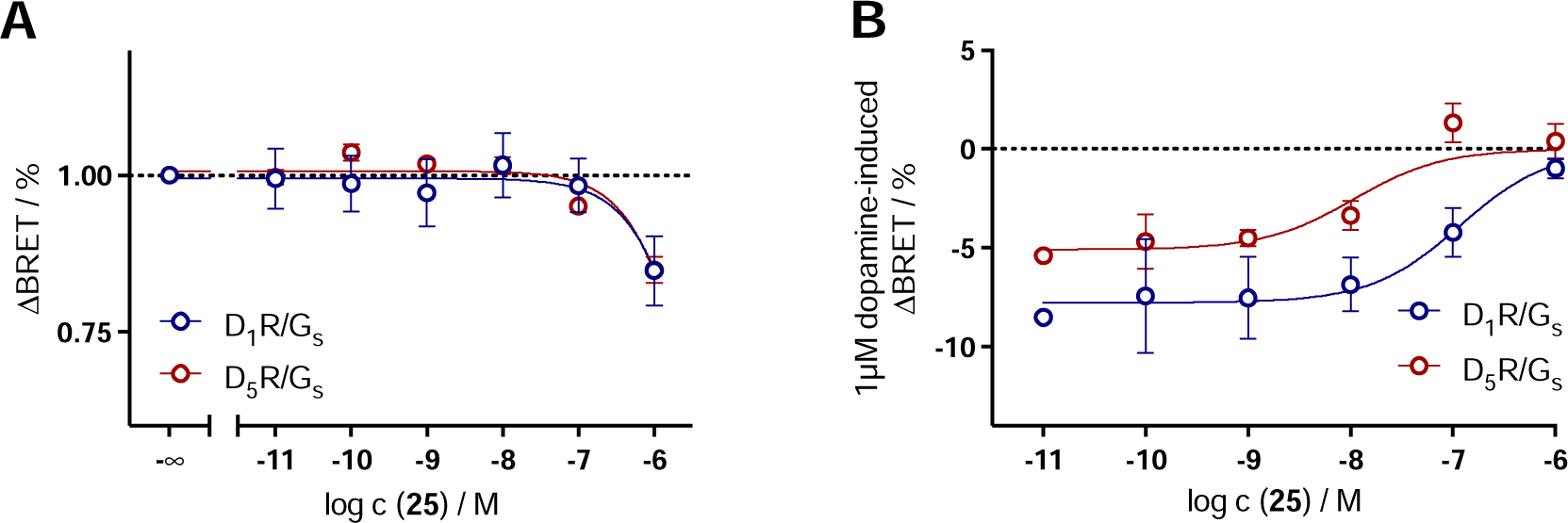
Concentration−response curves (CRCs) for G_s_ activation of **25** in the absence (A) and presence (B) of 1 µM dopamine in HEK293A cells transiently expressing the G_s_ BRET sensor along with the wild-type D_1_R or D_5_R. Graphs represent the means of three independent experiments each performed in duplicates. Data were analyzed by nonlinear regression and were best fitted to sigmoidal concentration-response curves.

### Fluorescence Properties

The fluorescence properties are usually not heavily affected by the addition of a pharmacophore, if the fluorophore is chemically not changed, as it’s the case for the 5-TAMRA and DY549-P1 dye. Nevertheless, the absorption and emission spectra and the quantum yield should be determined for new fluorescent ligands. The knowledge of the excitation and emission spectra and a high quantum yield are essential for their use in pharmacological assays and fluorescence microscopy. The emission and excitation spectra of **25**-**28** were recorded in PBS buffer containing 1 % of BSA (Figure 4). The excitation and emission maxima are presented in Table 3.

**Figure 4.**
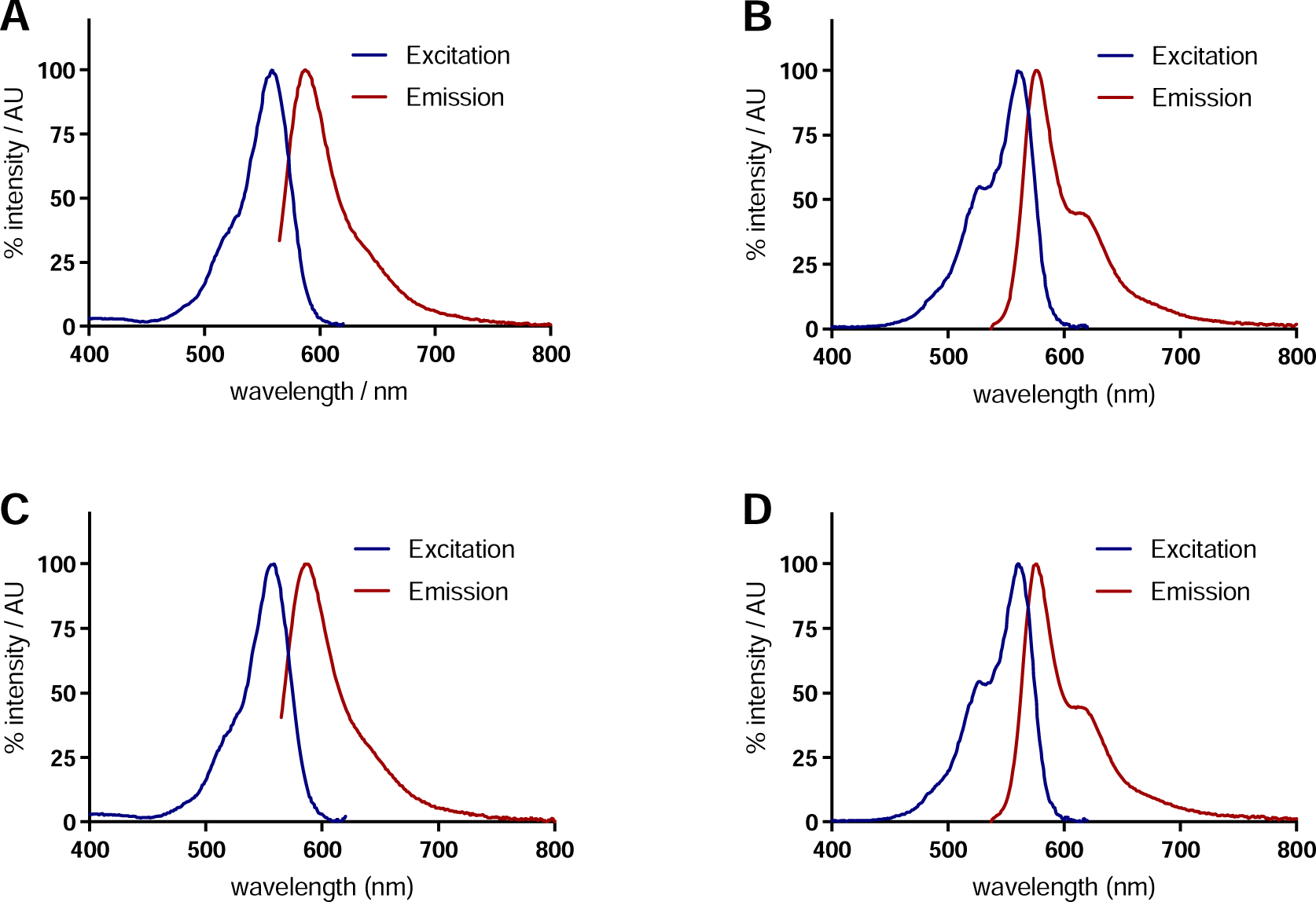
Excitation and emission spectra of **25** (A), **26** (B), **27** (C), and **28** (D) recorded in PBS buffer containing 1 % of BSA. The excitation wavelength for the emission spectra was 555 nm (**25**, **27**) or 527 nm (**26**, **28**). The emission wavelength collected during the excitation scan was 630 nm.

**Table 3.**
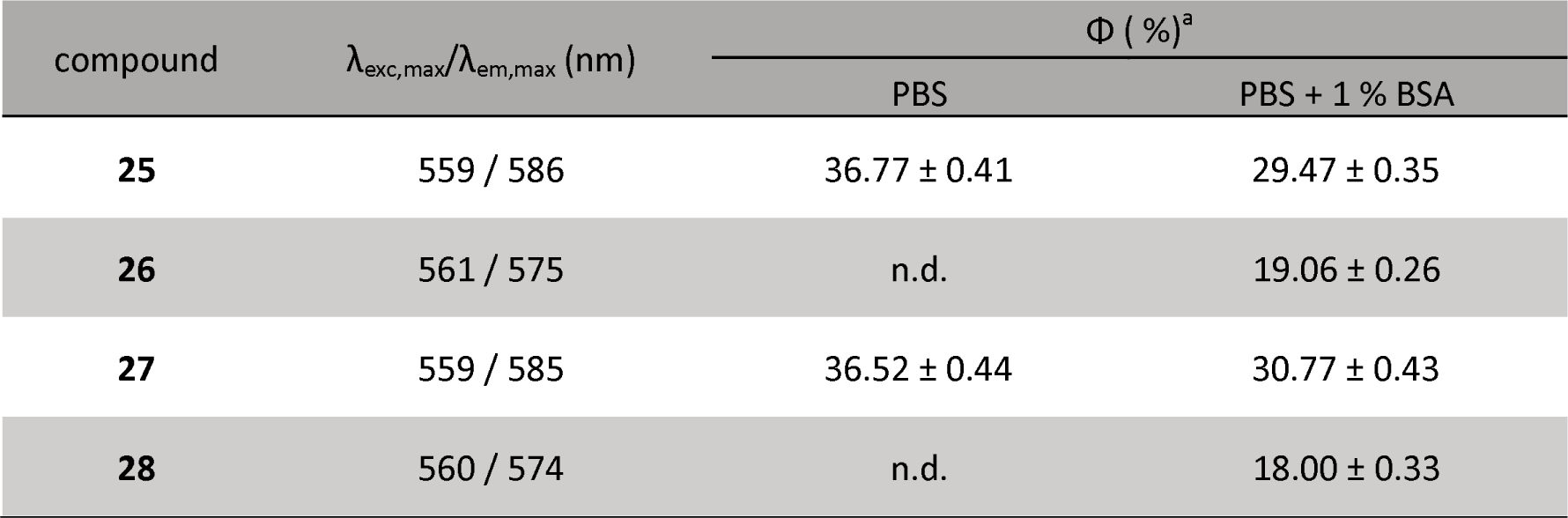
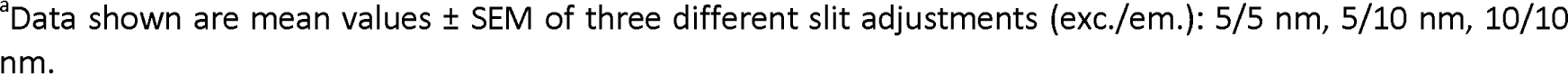
Fluorescence properties of compounds **25-28**.

| compound | $\lambda_{\text{exc,max}}/\lambda_{\text{em,max}}$ (nm) | $\Phi$ (%)<sup>a</sup> | |
| --- | --- | --- | --- |
|  |  | PBS | PBS + 1 % BSA |
| **25** | 559 / 586 | 36.77 ± 0.41 | 29.47 ± 0.35 |
| **26** | 561 / 575 | n.d. | 19.06 ± 0.26 |
| **27** | 559 / 585 | 36.52 ± 0.44 | 30.77 ± 0.43 |
| **28** | 560 / 574 | n.d. | 18.00 ± 0.33 |

<sup>a</sup>Data shown are mean values ± SEM of three different slit adjustments (exc./em.): 5/5 nm, 5/10 nm, 10/10 nm.

The quantum yield of the compounds were determined in PBS buffer containing 1 % BSA for all four compounds and additionally in PBS buffer for compounds **25** and **27** (Table 3) following a previously published protocol.[52] The 5-TAMRA-labeled compounds **25** and **27** have a higher quantum yield of approximately 36 % in PBS buffer compared to the DY549-P1-labeled ligands (**26** and **28**; approx. 19 %). Furthermore, it was observed that the addition of 1 % BSA led to a decrease in the quantum yield of the 5-TAMRA-labeled fluorescent ligands, but still with a satisfactory quantum yield of approx. 30 %. Based on these results all four ligands, but especially **25** and **27**, should be suitable for the use in fluorescence microscopy.

### Microscopy

In order to test the suitability of **25** for laser scanning confocal microscopy (LSCM) and to visualize the binding of **25** to the D_1_R, HEK-293T cells were transiently transfected with the D_1_R C-terminally tagged with YFP. 48 hours after transfection confocal microscopy images were recorded (Figure S8, SI). After addition of 50 nM of **25** a rapid accumulation of fluorescence at the cell surface was observed. This is caused by the fast association of **25** to the D_1_R expressed on the cell membrane reaching saturation binding in less than one minute. Based on the fluorescence intensity of individual cells, a time-dependent association curve could be obtained illustrating the fast binding of **25** to the D_1_R followed by a constant dissociation of **25** from the receptor after addition of 50 µM SCH-23390 (1,000-fold excess) (Figure 5). Based on the association and dissociation rates a kinetic p*K_d_* value of 8.07 was calculated (Table 4). These results demonstrate the suitability of **25** for the use in fluorescence microscopy as a labeling agent to visualize the D_1_R in live cells.

**Figure 5.**
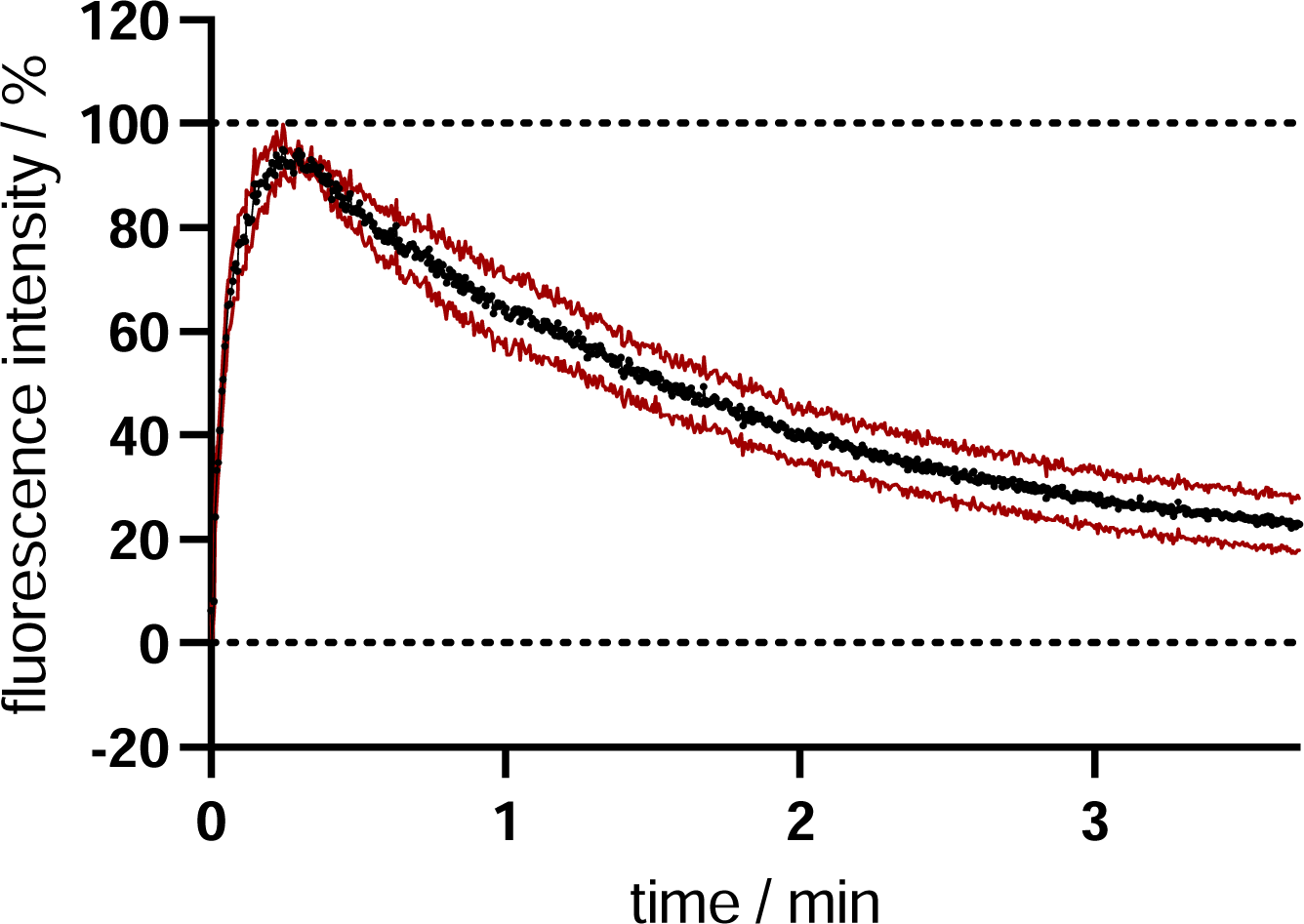
Binding kinetics of **25** at the hD_1_R in whole HEK-293T cells using LSCM. Graph shows association of **25** (c = 50 nM) to the receptor and dissociation of **25** induced by addition of SCH-23390 (c = 50 µM, 1,000-fold excess) from a representative experiment. Data represent mean ± SEM of four independent cells of one experiment.

**Table 4.**
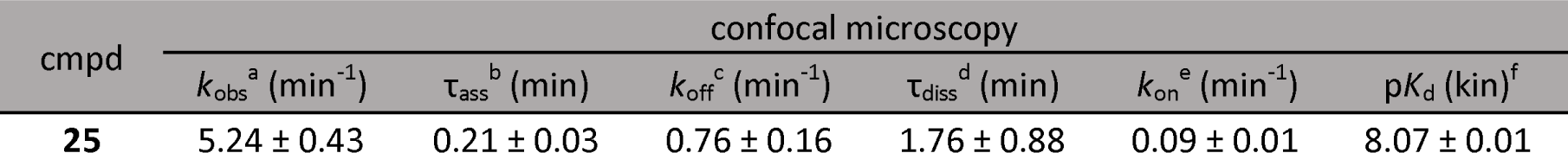
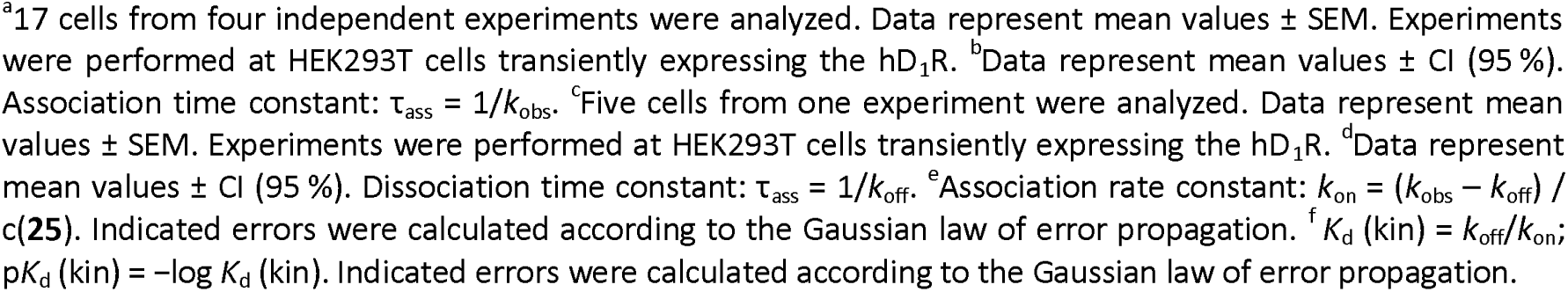
Kinetic binding constants of **25** at the hD_1_R in confocal microscopy.

### Molecular Brightness

To further characterize the ligand’s binding affinity in intact, living cells we employed molecular brightness to calculate the number of receptors decorated by the ligand on the basal membrane of cells, for varying ligand concentrations.[53] In order to reference the fraction of ligand-receptor complexes to the total amount of receptors expressed, we transfected HEK293-AD cells with D_1_R C-terminally tagged with the photostable fluorescent protein mNeonGreen (D1R-mNeonGreen). Cells were incubated with increasing concentrations of **25** and after reaching equilibrium, ligand was washed out and cell’s basal membranes were imaged over time in the two spectral channels (GFP /receptor and TAMRA/ligand) using a confocal microscope (Figure 6A). From the obtained movies the average number of emitters within the confocal excitation volume were calculated based on the fluctuation of the fluorescence photon counts within each pixel (see methods) and plotted against each other. A linear regression was fitted to the data points and the slope calculated (Figure 6B). A slope of 1 signifies a 1:1 ratio in the number of GFPs and TAMRAs and with this a full occupancy of the receptor by the ligand. Slopes between 0 and 1 represent partial occupancy. Plotting obtained slopes versus logarithmic expression of ligand concentration returns a sigmoid concentration response curve and yields a pK_d_ of 8.92 ± 0.13, that matches well the results obtained by radioligand binding (Figure 6C).

**Figure 6.**
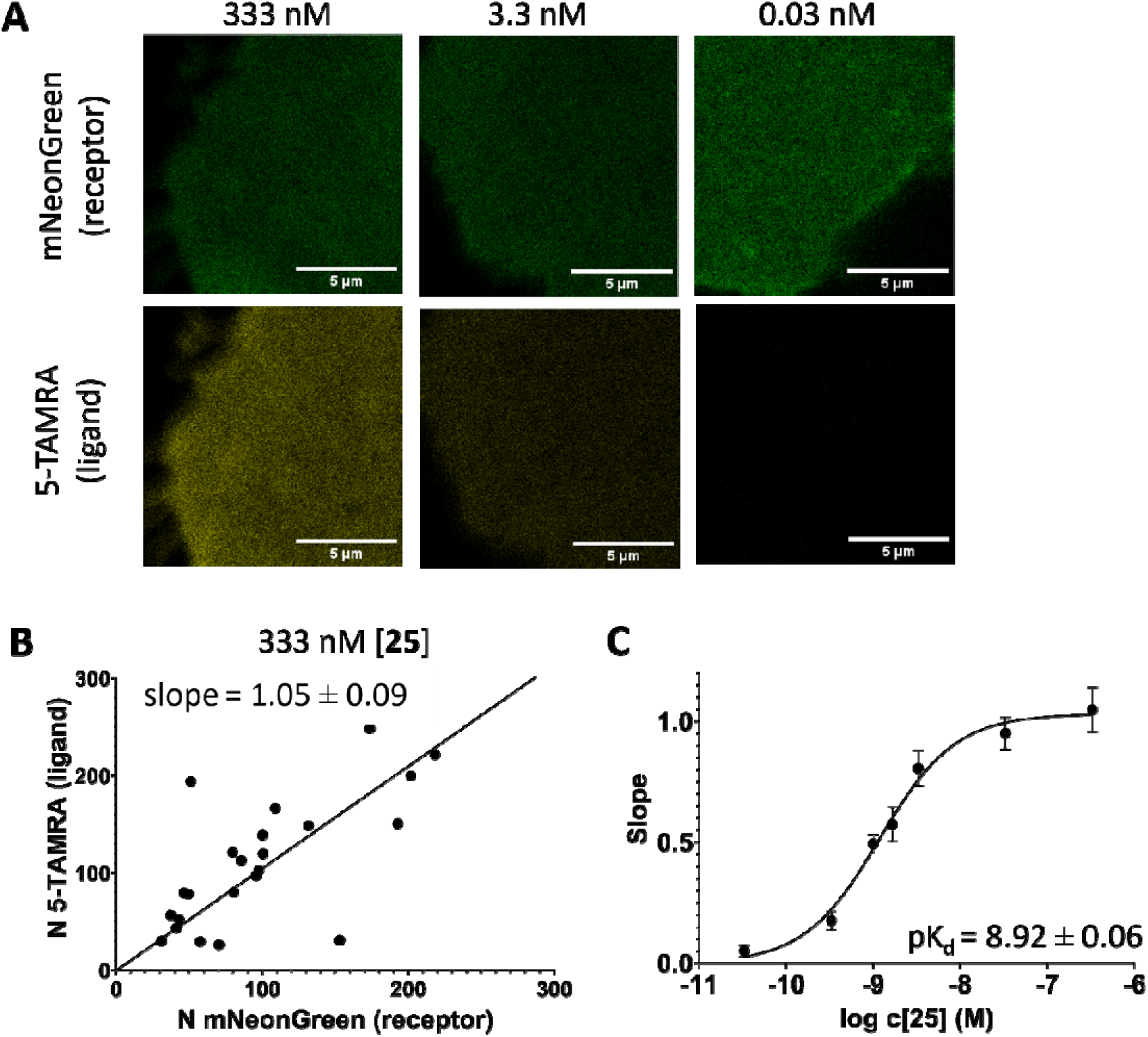
Association of **25** to the hD_1_R using molecular brightness analysis. Basolateral membranes of HEK-293AD cells transiently expressing hD_1_R-mNeonGreen and preincubated with indicated amount of **25** (**A**); calculated numbers of emitters for 5-TAMRA and mNeonGreen channels, and the corresponding linear regression fit (mean ± SEM; n = 23 cells from 3 independent experiments) (**B**), slopes obtained from linear fits plotted against log concentrations of **25** with the corresponding non-linear fit and pK_d_ value (mean ± SEM; n = at least 20 cells from 3 independent experiments for each datapoint) (**C**).

## Conclusion

A set of six different fluorescent ligands containing linkers of different lengths and chemical compositions and two different fluorescent dyes was designed and synthesized. The known D_1_R/D_5_R antagonistic SCH-23390 was used as a pharmacophore for the fluorescent ligands. Two fluorescent dyes, 5-TAMRA and DY549-P1, were chosen because of their proposed suitability as a fluorescent tracer for microscopy studies. After the successful synthesis and purification of the ligands, their fluorescent and pharmacological properties were determined. Radioligand binding studies were performed revealing **25** and **27** as D_1_-like receptor selective fluorescent ligands with binding affinity in the low nanomolar range. Compound **25**, equipped with the medium-length hydrophilic PEG linker and 5-TAMRA as dye, showed the best overall results regarding affinity, selectivity, quantum yield and was used as an exemplary compound to determine its mode of action. A G protein biosensor assay confirmed the expected neutral antagonism of **25**. Furthermore, **25** was used successfully to label D_1_Rs in live cells for LSCM demonstrating its suitability for fluorescence microscopy. In molecular brightness studies, the ligand’s binding affinity could be determined in a range that was in good agreement with radioligand binding data and a full occupancy of the receptor by the ligand. Overall, the set of fluorescent ligands, especially **25**, represent a versatile tool for different experimental setups for further investigations at the D_1_-like receptors.

## Experimental Section

### Chemistry

Commercially available chemicals and solvents were purchased from standard commercial suppliers (Merck (Darmstadt, Germany), Sigma-Aldrich (Munich, Germany), Acros Organics (Geel, Belgium), Alfa Aesar (Karlsruhe, Germany), abcr (Karlsruhe, Germany) or TCI Europe (Zwijndrecht, Belgium) and were used as received. All solvents were of analytical grade. The fluorescent dye 5-TAMRA NHS ester was purchased from Lumiprobe (Hannover, Germany). The fluorescent dye DY549-P1 was purchased from Dyomics GmbH (Jena, Germany). Deuterated solvents for nuclear magnetic resonance (^1^H NMR and ^13^C NMR) spectra were purchased from Deutero GmbH (Kastellaun, Germany). All reactions carried out with dry solvents were accomplished in dry flasks under nitrogen or argon atmosphere. For the preparation of buffers, HPLC eluents, and stock solutions millipore water was used. Column chromatography was accomplished using Merck silica gel Geduran 60 (0.063-0.200 mm). Flash chromatography was performed using an Agilent Technologies 971-FP Flash Purification System (Agilent Technologies, Santa Clara, CA) with SI-HP 30 µm puriFlash columns from Interchim (Montlucon, France). The reactions were monitored by thin layer chromatography (TLC) on Merck silica gel 60 F254 aluminium sheets and spots were visualized under UV light at 254 nm, by potassium permanganate or ninhydrin staining. Lyophilization was done with a Christ alpha 2-4 LD equipped with a vacuubrand RZ 6 rotary vane vacuum pump. Nuclear magnetic resonance (^1^H NMR and ^13^C NMR) spectra were recorded on a Bruker (Karlsruhe, Germany) Avance 300 (^1^H: 300 MHz, ^13^C: 75 MHz), 400 (^1^H: 400 MHz, ^13^C: 101 MHz) or 600 (^1^H: 600 MHz) spectrometer using perdeuterated solvents. The chemical shift δ is given in parts per million (ppm). Multiplicities were specified with the following abbreviations: s (singlet), d (doublet), t (triplet), q (quartet), quin (quintet), m (multiplet), and br (broad signal) as well as combinations thereof. ^13^C NMR-Peaks were determined by DEPT 135 and DEPT 90 (distortionless enhancement by polarization transfer). NMR spectra were processed with MestReNova 11.0 (Mestrelab Research, Compostela, Spain). High-resolution mass spectrometry (HRMS) was performed on an Agilent 6540 UHD Accurate-Mass Q-TOF LC/MS system (Agilent Technologies, Santa Clara, CA) using an ESI source. Preparative HPLC was performed with a system from Waters (Milford, Massachusetts, USA) consisting of a 2524 binary gradient module, a 2489 detector, a prep inject injector and a fraction collector III. A Phenomenex Gemini 5µm NX-C18 column (110Å, 250 x 21.2mm, Phenomenex Ltd., Aschaffenburg, Germany) served as stationary phase. As mobile phase, 0.1% TFA or 0.1% NH_3_ in millipore water and acetonitrile (MeCN) were used. The temperature was 25 °C, the flow rate 20 mL/min and UV detection was performed at 220 nm. Analytical HPLC experiments were performed on a 1100 HPLC system from Agilent Technologies equipped with Instant Pilot controller, a G1312A Bin Pump, a G1329A ALS autosampler, a G1379A vacuum degasser, a G1316A column compartment and a G1315B DAD detector. The column was a Phenomenex Gemini 5µm NX-C18 column (110Å, 250 x 4.6mm, Phenomenex Ltd., Aschaffenburg, Germany) tempered at 30 °C. As mobile phase, mixtures of MeCN and aqueous TFA (for compounds **23**, **25**, **27**) or aqueous NH_3_ (for compounds **24**, **26**, **28**) were used (linear gradient: MeCN/TFA (0.1%) (v/v) 0 min: 10:90, 25-35 min: 95:5, 36-45 min: 10:90; flow rate = 1.00 mL/min, t_0_ = 3.21 min).

Capacity factors were calculated according to k = (t_R_ - t_0_)/t_0_. Detection was performed at 220 nm. Furthermore, a filtration of the stock solutions with PTFE filters (25 mm, 0.2 µm, Phenomenex Ltd., Aschaffenburg, Germany) was carried out before testing. Compound purities determined by HPLC were calculated as the peak area of the analyzed compound in % relative to the total peak area (UV detection at 220 nm). The HPLC purity and stability of final compounds is displayed in the SI (cf. Figure S1-S6, SI).

## Synthesis and Analytical Data

### 2-Bromo-1-(4-nitrophenyl)ethan-1-one (1)[54]

4-Nitroacetophenone (10.00 g, 60.54 mmol, 1 eq.) was dissolved in DCM and added to a suspension of ***N***-bromosuccinimide (12.92 g, 72.64 mmol, 1.2 eq.) and *p*-toluenesulfonic acid (1.14 g, 6.06 mmol, 0.1 eq.) in DCM at room temperature. The reaction was heated to reflux overnight. The reaction was poured on water and the organic phase was separated. The aqueous phase was extracted with DCM, the combined organic phases washed with saturated bicarbonate solution and brine, dried over Na_2_SO_4_ and the solvent was removed under reduced pressure. The crude product was purified by column chromatography (DCM/PE 1:1). A pale yellow solid was obtained (9.41 g, 64%). R_f_ = 0.61 (PE/DCM 4:6). ^1^H NMR (300 MHz, CDCl_3_) δ 8.48 – 8.26 (m, 1H), 8.21 – 8.07 (m, 1H), 4.47 (s, 1H). ^13^C NMR (75 MHz, CDCl_3_) δ 189.92, 150.71, 138.37, 130.11, 124.08, 30.18. HRMS (EI-MS): m/z M^ill+^ calculated for C_8_H_6_NO_3_Br^ill+^: 242.9526, found 242.9522; C_8_H_6_NO_3_Br (244.04).

### 2-Bromo-1-(4-nitrophenyl)ethan-1-ol (2)[55]

*1* (3.34 g, 13.70 mmol, 1 eq.) was dissolved in MeOH. NaBH_4_ (0.18 g, 4.80 mmol, 0.35 eq.) was added in portions at 0°C. The reaction was stirred for 1 h at room temperature. The solvent was removed under reduced pressure and the crude product was dissolved in water. The aqueous phase was extracted with diethyl ether. The combined organic phases were washed with saturated ammonium chloride solution and brine, dried over Na_2_SO_4_ and the solvent was removed under reduced pressure. A slightly yellow solid was obtained (2.23 g, 66%). The product was used without further purification.

### 2-(4-*N*itrophenyl)oxirane (3)[56]

*2* (0.5 g, 2.03 mmol, 1 eq.) and K_2_CO_3_ (0.56 g, 4.06 mmol, 2 eq.) were dissolved in THF and heated to reflux overnight. The solid was filtered off and the solvent was removed under reduced pressure. The crude product was purified by column chromatography (PE/EtOAc 8:2). A yellow solid was obtained (0.29 g, 87%). R_f_ = 0.78 (PE/EtOAc 7:3). ^1^H NMR (300 MHz, CDCl_3_) δ 8.29 – 8.13 (m, 2H), 7.56 – 7.36 (m, 2H), 3.96 (dd, *J* = 4.1, 2.5 Hz, 1H), 3.23 (dd, *J* = 5.5, 4.1 Hz, 1H), 2.78 (dd, *J* = 5.5, 2.5 Hz, 1H). ^13^C NMR (75 MHz, CDCl_3_) δ 145.26, 126.24, 123.85, 51.71, 51.48. HRMS (EI-MS): m/z M^ill+^ calculated for C_8_H_7_NO_3_^ill+^: 164.0342, found 164.0339; C_8_H_7_NO_3_ (165.15).

### 3-Chloro-4-methoxybenzaldehyde (4)[57]

Sulfurylchloride (2.41 ml, 29.38 mmol, 2 eq.) was added to a solution of 4-methoxybenzaldehyde (2.00 g, 14.69 mmol, 1 eq.) in acetic acid. The reaction was stirred overnight at room temperature and poured onto a mixture of ice and water. The white solid was filtered off, washed with ice cold water and hexanes and dried *in vacuo*. A white solid was obtained (2,15 g, 86%). R_f_ = 0.68 (PE/EtOAc 7:3). 1H NMR (300 MHz, CDCl3) δ 9.85 (s, 1H), 7.91 (d, *J* = 2.0 Hz, 1H), 7.78 (dd, *J* = 8.5, 2.0 Hz, 1H), 7.05 (d, *J* = 8.5 Hz, 1H), 3.99 (s, 3H). 13C NMR (75 MHz, CDCl3) δ 189.8, 159.9, 131.3, 130.6, 130.3, 123.7, 111.7, 56.5. HRMS (APCI-MS): m/z [M+H]^+^ calculated for C_8_H_8_ClO_2_^+^: 171.0207, found 171.0207; C_8_H_7_ClO_2_ (170.59).

### Chloro-1-methoxy-4-(2-nitrovinyl)benzene (5)[58]

*4* (6.40 g, 37.52 mmol, 1 eq.), nitromethane (6.03 ml, 112.56 mmol, 3 eq.) and ammonium acetate (2.40 g, 93.80 mmol, 2.5 eq.) were dissolved in acetic acid and heated to reflux for 4 h. Water was added and the reaction was extracted with DCM. The combined organic phases were washed with 1 N aq. NaOH solution and brine, dried over Na_2_SO_4_ and the solvent removed under reduced pressure. The crude product was purified via column chromatography (PE/EtOAc 8:2). A yellow solid was obtained (5.49 g, 68%). R_f_ = 0.67 (PE/EtOAc 7:3). ^1^H NMR (300 MHz, CDCl_3_) δ 7.90 (d, *J* = 13.6 Hz, 1H), 7.58 (d, *J* = 2.2 Hz, 1H), 7.50 (d, *J* = 13.6 Hz, 1H), 7.44 (dd, *J* = 8.7, 2.3 Hz, 1H), 6.98 (d, *J* = 8.6 Hz, 1H), 97 (s, 3H). ^13^C NMR (75 MHz, CDCl_3_) δ 158.08, 137.69, 136.02, 130.48, 129.83, 123.83, 123.33, 112.36, 56.46. HRMS (EI-MS): m/z M^ill+^ calculated for C_9_H_8_ClNO_3_^ill+^: 213.0180, found 213.0187; C_9_H_8_ClNO_3_ (213.67).

### 2-(3-Chloro-4-methoxyphenyl)ethan-1-amine (6)[58]

*5* (5.43 g, 25.42 mmol, 1 eq.) was dissolved in THF and added slowly at room temperature to a suspension of LiAlH_4_ (2.89 g, 76.26 mmol, 3 eq.) in THF. The reaction was heated to reflux for 3 h. After cooling to room temperature water (10 ml) and 20% aq. KOH-solution were added carefully at 0°C and the reaction was stirred for 30 min at room temperature. The white solid was filtered off and the filtrate was dried under reduced pressure. The crude product was dried *in vacuo*. A yellow oil was obtained (4.44 g, 94%). R_f_ = 0.05 (DCM/MeOH 98:2). ^1^H NMR (300 MHz, CDCl_3_) δ 7.20 (d, *J* = 2.1 Hz, 1H), 7.05 (dd, *J* = 8.3, 2.2 Hz, 1H), 6.86 (d, *J* = 8.4 Hz, 1H), 3.87 (s, 3H), 2.92 (t, *J* = 6.8 Hz, 2H), 2.66 (t, *J* = 6.8 Hz, 2H), 1.49 (bs, 2H). ^13^C NMR (75 MHz, CDCl_3_) δ 153.45, 132.98, 130.47, 128.03, 122.26, 112.13, 56.20, 43.46, 38.78. HRMS (ESI-MS): m/z [M+H]^+^ calculated for C_9_H_13_ClNO^+^: 186.0680, found 186.0679; C_9_H_12_ClNO (185,65).

### tert-Butyl (3-Chloro-4-methoxyphenethyl)carbamate (7)

Di-*tert*-butyldicarbonate (3.23 g, 14.82 mmol, 1.1 eq.) was dissolved in DCM and added slowly at room temperature to a solution of **6** (2.50 g, 13.47 mmol, 1 eq.) in DCM. The reaction was stirred at room temperature overnight. The solvent was removed under reduced pressure and the crude product was dried *in vacuo*. A slightly yellow solid was obtained (3.79 g, 98%). R_f_ = 0.48 (PE/EtOAc 8:2). ^1^H NMR (400 MHz, CDCl_3_) δ 7.19 (d, *J* = 2.1 Hz, 1H), 7.03 (dd, *J* = 8.4, 2.1 Hz, 1H), 6.85 (d, *J* = 8.4 Hz, 1H), 3.87 (s, 3H), 3.31 (t, *J* = 7.0 Hz, 2H), 2.70 (t, *J* = 7.0 Hz, 2H), 1.42 (s, 9H). ^13^C NMR (101 MHz, CDCl_3_) δ 155.90, 153.61, 132.15, 130.49, 127.98, 122.33, 112.22, 56.18, 40.89, 35.11, 28.39, 23.86. HRMS (ESI-MS): m/z [M+H]^+^ calculated for C_14_H_20_ClNO_3_^+^: 286.1204, found 286.1204; C_14_H_20_ClNO_3_ (285.77).

### 2-(3-Chloro-4-methoxyphenyl)-*N*-methylethan-1-amine (8)[59]

*7* (6.2 g, 21.70 mmol, 1 eq.) was dissolved in THF and added slowly at room temperature to a suspension of LiAlH_4_ (2.47 g, 65.10 mmol, 3 eq.) in THF. The reaction was heated to reflux for 3 h. After cooling to room temperature water (10 ml) and 20% aq. KOH-solution were added carefully at 0°C and the reaction was stirred for 30 min at room temperature. The white solid was filtered off and the filtrate was dried under reduced pressure. The crude product was dried *in vacuo*. A yellow oil was obtained (3.88 g, 90%). R_f_ = 0.16 (DCM/MeOH+1%NH_3_ 95:5). ^1^H NMR (300 MHz, CDCl_3_) δ 7.21 (d, *J* = 2.1 Hz, 1H), 7.05 (dd, *J* = 8.4, 2.2 Hz, 1H), 6.85 (d, *J* = 8.4 Hz, 1H), 3.87 (s, *J* = 3.9 Hz, 3H), 2.84 – 2.76 (m, 2H), 2.76 – 2.68 (m, 2H), 2.43 (s, 3H). ^13^C NMR (75 MHz, CDCl_3_) δ 153.42, 133.24, 130.34, 127.92, 122.27, 112.13, 77.48, 77.06, 76.63, 56.20, 53.13, 36.38, 35.03. HRMS (ESI-MS): m/z [M+H]^+^ calculated for C_10_H_15_ClNO^+^: 200.0837, found 200.0839; C_10_H_14_ClNO (199.69).

### 2-((3-Chloro-4-methoxyphenethyl)(methyl)amino)-1-(4-nitrophenyl)ethan-1-ol (9)[48]

*8* (2.44 g, 12.22 mmol, 1 eq.) and *3* (2.02 g, 12.22 mmol, 1 eq.) were dissolved in acetonitrile and heated to reflux overnight. The solvent was removed under reduced pressure and the crude product was purified by column chromatography (DCM/MeOH+1%NH_3_ 98:2). A brown oil was obtained (4.00 g, 90%). R_f_ = 0.16 (DCM/MeOH+1%NH_3_ 98:2). ^1^H NMR (300 MHz, CDCl_3_) δ 8.23 – 8.12 (m, 2H), 7.57 – 7.47 (m, 2H), 7.20 (d, *J* = 2.2 Hz, 1H), 7.05 (dd, *J* = 8.4, 2.2 Hz, 1H), 6.86 (d, *J* = 8.4 Hz, 1H), 4.78 (dd, *J* = 10.4, 3.5 Hz, 1H), 4.19 (bs, 1H), 3.87 (s, 3H), 2.90 – 2.57 (m, 5H), 2.52 – 2.41 (m, 4H). ^13^C NMR (75 MHz, CDCl_3_) δ 153.54, 149.73, 147.33, 132.59, 130.34, 127.85, 126.52, 123.64, 122.32, 112.16, 68.56, 65.26, 59.18, 56.19, 41.73, 32.47. HRMS (ESI-MS): m/z [M+H]^+^ calculated for C_18_H_22_ ClN_2_O_4_^+^: 365.1263, found 365.1268; C_18_H_21_ClN_2_O_4_ (364.83).

### Chloro-8-methoxy-3-methyl-1-(4-nitrophenyl)-2,3,4,5-tetrahydro-1*H*-benzo[d]azepine (10)

*9* (2.49 g, 7.65 mmol, 1 eq.) was dissolved in Eaton’s reagent (40 ml) and stirred for 72 h at room temperature. The reaction was poured onto ice water and basified with 20% aqueous KOH-solution. The aqueous phase was extracted with DCM. The combined organic phases were washed with water, dried over Na_2_SO_4_ and the solvent was removed under reduced pressure. The crude product was purified by column chromatography (DCM/MeOH+1% NH_3_ 98:2). A red solid was obtained (1.20 g, 45%). R_f_ = 0.18 (DCM/MeOH+1%NH_3_ 98:2). ^1^H NMR (300 MHz, CDCl_3_) δ 8.22 – 8.10 (m, 2H), 7.37 – 7.28 (m, 2H), 7.14 (s, 1H), 6.39 (s, 1H), 4.42 – 4.27 (m, 1H), 3.71 (s, 3H), 3.26 – 3.13 (m, 1H), 2.96 – 2.75 (m, 2H), 2.73 – 2.52 (m, 3H), 2.38 (s, 3H).^13^C NMR (75 MHz, CDCl_3_) δ 153.29, 149.94, 146.48, 141.87, 134.37, 131.64, 129.06, 123.72, 120.27, 113.44, 61.34, 57.29, 56.15, 50.27, 47.93, 34.98. HRMS (ESI-MS): m/z [M+H]^+^ calculated for C_18_H_20_ClN_2_O_3_^+^: 347.1157, found 347.1162; C_18_H_19_ClN_2_O_3_ (346.81).

### Chloro-3-methyl-5-(4-nitrophenyl)-2,3,4,5-tetrahydro-1*H*-benzo[d]azepin-7-ol (11)[48]

*10* (0.80 g, 2.31 mmol, 1 eq.) was dissolved DCM and cooled to -78°C under Ar-atmosphere. BBr_3_ (0.33 ml, 3.45 mmol, 1.5 eq.) dissolved in DCM was added at -78°C and the reaction stirred for 1 h at this temperature and after that stirred at room temperature overnight. MeOH (5 ml) was added at - 78°C and the reaction was stirred for 1 h at room temperature. The solvent was removed under reduced pressure and the product was dried *in vacuo*. A brown solid was obtained (0.80 g, 84%). The product was used without further purification. HRMS (ESI-MS): m/z [M+H]^+^ calculated for C_17_H_18_ClN_2_O_3_^+^: 333.1000, found 333.1004; C_17_H_17_ClN_2_O_3_*HBr (413.70).

### 8-Chloro-3-methyl-5-(4-nitrophenyl)-2,3,4,5-tetrahydro-1*H*-benzo[d]azepin-7-yl acetate (12a)

*11* (0.13 g, 0.35 mmol, 1 eq.) and NEt_3_ (98.4 µl, 0.70 mmol, 2 eq.) were dissolved in DCM and cooled to 0°C. Acetyl chloride (37.8 µl, 0.53 mmol, 1.5 eq.) was added slowly and the reaction was stirred at room temperature overnight. The solvent was removed under reduced pressure and the product was purified by column chromatography (98:2 DCM/MeOH). An orange solid was obtained (0.10 g, 77%). R_f_ = 0.48 (98:2 DCM/MeOH+1%NH_3_). ^1^H NMR (300 MHz, CDCl_3_) δ 8.23 – 8.17 (m, 2H), 7.37 – 7.30 (m, 2H), 7.24 (s, 1H), 6.41 (s, 1H), 4.37 (d, *J* = 7.5 Hz, 1H), 3.09 – 2.91 (m, 3H), 2.81 – 2.73 (m, 2H), 2.54 – 2.45 (m, 1H), 2.40 (s, 3H), 2.27 (s, 3H). ^13^C NMR (75 MHz, CDCl_3_) δ 168.52, 149.57, 146.74, 145.15, 142.67, 140.44, 131.40, 129.21, 124.78, 123.98, 123.52, 61.67, 56.66, 49.53, 47.77, 35.57, 20.62. HRMS (ESI-MS): m/z [M+H]^+^ calculated for C_19_H_20_ClN_2_O_4_^+^: 375.1106, found 375.1113; C_19_H_19_ClN_2_O_4_ (374.82).

### Chloro-3-methyl-1-(4-nitrophenyl)-8-((triisopropylsilyl)oxy)-2,3,4,5-tetrahydro-1*H*-benzo[d]azepine (12b)

Triisopropylsilylchloride (309 µl, 1.46 mmol, 2 eq.) and imidazole (0.10g, 1.46 mmol, 2 eq.) were dissolved in DMF. *11* (0.30 g, 0.73 mmol, 1 eq.) and NEt_3_ (513 µl, 3.65 mmol, 5 eq.) dissolved in DMF were added at room temperature under Ar-atmosphere. The reaction was stirred overnight at room temperature and subsequently poured onto water. The aqueous phase was extracted with diethyl ether. The combined organic phases were washed with saturated ammonium chloride solution and brine, dried over Na_2_SO_4_ and the solvent removed under reduced pressure. The crude product was purified by flash chromatography (SiO_2_, 0 min to 20 min, 100:0 to 95:5 DCM/MeOH). R_f_ = 0.40 (DCM/MeOH 95:5). A slightly yellow solid was obtained (0.32 g, 90%). ^1^H NMR (300 MHz, CDCl_3_) δ 8.29 – 8.15 (m, 2H), 7.38 – 7.28 (m, 1H), 7.12 (s, 1H), 6.06 (s, 1H), 4.53 – 4.28 (m, 1H), 3.13 – 2.83 (m, 4H), 2.78 – 2.63 (m, 1H), 2.50 – 2.35 (m, 4H), 1.05 – 0.88 (m, 21H). ^13^C NMR (75 MHz, CDCl_3_) δ 150.35, 150.15, 146.69, 142.27, 134.19, 131.25, 129.22, 123.85, 122.65, 120.20, 61.91, 57.25, 49.02, 47.66, 34.92, 17.77, 12.71. HRMS (ESI-MS): m/z [M+H]^+^ calculated for C_26_H_38_ClN_2_O_3_Si^+^: 489.2335, found 489.2342; C_26_H_37_ClN_2_O_3_Si (488.23).

### 5-(4-Aminophenyl)-8-chloro-3-methyl-2,3,4,5-tetrahydro-1*H*-benzo[d]azepin-7-yl acetate (13a)

*12a* (0.14 g, 0.37 mmol, 1 eq.) and Pd/C (10%) (0.01 g, 10 wt%) were dissolved in a mixture of THF and MeOH (1:1) and the reaction was stirred overnight under H_2_-atmosphere. The reaction was filtered over celite and the solvent was removed under reduced pressure. The crude product was dried *in vacuo.* A yellow solid was obtained (0.12 g, 95%). The product was used without further purification. HRMS (ESI-MS): m/z [M+H]^+^ calculated for C_19_H_22_ClN_2_O_2_^+^: 345.1364, found 345.1370; C_19_H_21_ClN_2_O_2_ (344.84).

### 4-(7-Chloro-3-methyl-8-((triisopropylsilyl)oxy)-2,3,4,5-tetrahydro-1*H*-benzo[d]azepin-1-yl)aniline (13b)

*12b* (0.42 g, 0.86 mmol, 1 eq.) and Pd/C (10%) (0.04 g, 10 wt%) were dissolved in a mixture of THF and MeOH (1:1) and the reaction was stirred overnight under H_2_-atmosphere. The reaction was filtered over celite and the solvent was removed under reduced pressure. The crude product was dried *in vacuo.* A brown oil was obtained (0.38 g, 96%). The product was used without further purification. HRMS (ESI-MS): m/z [M+H]^+^ calculated for C_26_H_40_ClN_2_OSi^+^: 459.2593, found 459.2598; C_26_H_39_ClN_2_OSi (459.15).

### 3-(1,3-Dioxoisoindolin-2-yl)propane-1-sulfonyl chloride (14a)[49]

Thiourea (0.38 g, 5.00 mmol, 1 eq.) and 3-bromopropylphthalimide (1.34 g, 5.00 mmol, 1 eq.) were dissolved in EtOH and heated to reflux for 1 h. The solvent was removed under reduced pressure and the resulting white solid was added to a suspension of *N*-chlorosuccinimide (NCS) in MeCN and 2 N aq. HCl (1.62 ml, 3.24 mmol, 0.65 eq.) at 10°C. The reaction was stirred at 10°C for 30 min and the reaction was quenched with water. The aqueous phase was extracted with diethyl ether. The organic phase was washed with brine, dried over Na_2_SO_4_ and the solvent was removed under reduced pressure. The crude product was purified by column chromatography (3:1 PE/EtOAc). A white solid was obtained (1.29 g, 89%). R_f_ = 0.80 (1:1 PE/EtOAc). ^1^H NMR (300 MHz, CDCl_3_) δ 7.90 – 7.82 (m, 2H), 7.79 – 7.72 (m, 2H), 3.89 (t, *J* = 6.5 Hz, 2H), 3.80 – 3.70 (m, 2H), 2.51 – 2.35 (m, 2H). ^13^C NMR (75 MHz, CDCl_3_) δ 168.20, 134.43, 131.74, 123.61, 62.86, 35.55, 24.13. HRMS (ESI-MS): m/z [M+H]^+^ calculated for C_11_H_11_ClNO_4_S^+^: 288.0092, found 288.0092; C_11_H_10_ClNO_4_S (287.71).

### *E*thyl 5-(chlorosulfonyl)pentanoate (14b)[49,60]

Ethyl 5-bromovalerate (3.00 g, 14.35 mmol, 1 eq.) and thiourea (1.08 g, 14.35 mmol) were dissolved in EtOH and heated to reflux overnight. The solvent was removed under reduced pressure and the obtained solid was added at 5°C to a suspension of NCS (9.58 g, 71.75 mmol, 5 eq.) in acetonitrile and 2 N aq. HCl (5 ml). The reaction was stirred for 20 min below 10°C and poured onto water. The aqueous phase was extracted with diethyl ether. The combined organic phases were washed with brine, dried over Na_2_SO_4_ and the solvent was removed under reduced pressure. The crude product was purified by column chromatography (PE/EtOAc 8:2). A clear oil was obtained (2.86 g, 87%). R_f_ = 0.45 (PE/EtOAc 8:2). ^1^H NMR (300 MHz, CDCl_3_) δ 4.14 (q, *J* = 7.1 Hz, 2H), 3.77 – 3.61 (m, 2H), 2.39 (t, *J* = 7.1 Hz, 2H), 2.16 – 1.98 (m, 2H), 1.91 – 1.74 (m, 2H), 1.30 – 1.21 (m, 3H). ^13^C NMR (75 MHz, CDCl_3_) δ 172.52, 64.96, 60.72, 33.32, 23.83, 22.84, 14.23. HRMS (ESI-MS): m/z [M+H]^+^ calculated for C_7_H_14_ClO_4_S^+^: 229.0296, found 229.0295; C_7_H_13_ClO_4_S (228.69).

### General procedure A

Ethylene glycol derivative (1 eq.) and NEt_3_ (2.5 eq.) were dissolved in DCM. Methanesulfonyl chloride (2 eq.) was added at 0°C and the reaction was stirred at room temperature overnight. The white solid was filtered off and the filtrate was washed with saturated bicarbonate solution and brine, dried over Na_2_SO_4_ and the solvent was removed under reduced pressure. The crude product was purified by column chromatography (DCM/MeOH 98:2).

### Oxybis(ethane-2,1-diyl) dimethanesulfonate (15a)[61]

The product was synthesized following general procedure A from diethylene glycol (5.00 g, 47.12 mmol, 1 eq.), methanesulfonyl chloride (7.29 ml, 94.24 mmol, 2 eq.) and NEt_3_ (16.42 ml, 117.80 mmol, 2.5 eq.). A yellow oil was obtained (11.87 g, 96 %). R_f_ = 0.80 (DCM/MeOH 98:2). ^1^H NMR (300 MHz, CDCl_3_) δ 4.37 – 4.28 (m, 4H), 3.78 – 3.68 (m, 4H), 3.02 (s, 6H). ^13^C NMR (75 MHz, CDCl_3_) δ 69.05, 68.97, 37.57. HRMS (ESI-MS): m/z [M+H]^+^ calculated for C_6_H_15_O_7_S_2_ ^+^: 263.0254, found 263.0254; C_6_H_14_O_7_S_2_ (262.29).

### (*E*thane-1,2-diylbis(oxy))bis(ethane-2,1-diyl) dimethanesulfonate (15b)[62]

The product was synthesized following general procedure A from triethylene glycol (7.00 g, 46.61 mmol, 1 eq.), methanesulfonyl chloride (7.21 ml, 93.22 mmol, 2 eq.) and NEt_3_ (16.24 ml, 116.53 mmol, 2.5 eq.). A yellow oil was obtained (13.48 g, 94 %). R_f_ = 0.80 (DCM/MeOH 98:2). ^1^H NMR (300 MHz, CDCl_3_) δ 4.38 – 4.28 (m, 4H), 3.75 – 3.69 (m, 4H), 3.63 (s, 4H), 3.03 (s, 6H). ^13^C NMR (75 MHz, CDCl_3_) δ 70.51, 69.25, 68.99, 37.62. HRMS (ESI-MS): m/z [M+H]^+^ calculated for C_8_H_19_O_8_S_2_^+^: 307.0515, found 307.0516; C_8_H_18_O_8_S_2_ (306.34).

### General procedure B

The di-mesylate product (1 eq.) and sodium azide (4 eq.) were dissolved in a mixture of EtOH and DMF (4:1) and the reaction was heated to reflux overnight. The solvents were removed under reduced pressure and the crude product was dissolved in diethyl ether. The organic phase was washed with saturated ammonium chloride solution and brine, dried over Na_2_SO_4_ and the solvent removed under reduced pressure. The product was dried *in vacuo*.

### 1-Azido-2-(2-azidoethoxy)ethane (16a)[63]

The product was synthesized following general procedure B from **15a** (5.00 g, 19.06 mmol, 1 eq.) and sodium azide (4.96 g, 76.24 mmol, 4 eq.). A yellow oil was obtained (2.91 g, 98%). R_f_ = 0.35 (DCM/MeOH 98:2). ^1^H NMR (300 MHz, CDCl_3_) δ 3.72 – 3.64 (m, 4H), 3.45 – 3.36 (m, 4H). ^13^C NMR (75 MHz, CDCl_3_) δ 70.10, 50.76. HRMS (APCI-MS): m/z [M+H]^+^ calculated for C_4_H_9_N_6_O^+^: 157.0832, found 157.0835; C_4_H_8_N_6_O (156.15).

### 1,2-Bis(2-azidoethoxy)ethane (16b)[64]

The product was synthesized following general procedure B from **15b** (6.00 g, 19.59 mmol, 1 eq.) and sodium azide (5.09 g, 78.36 mmol, 4 eq.). A yellow oil was obtained (3.97 g, 99%). R_f_ = 0.26 (DCM/MeOH 98:2). ^1^H NMR (300 MHz, CDCl_3_) δ 3.72 – 3.65 (m, 8H), 3.43 – 3.34 (m, 4H). ^13^C NMR (75 MHz, CDCl_3_) δ 70.76, 70.17, 50.71. HRMS (APCI-MS): m/z [M+H]^+^ calculated for C_6_H_13_N_6_O_2_^+^: 201.1095, found 201.1096; C_6_H_12_N_6_O_2_ (200.20).

### General procedure C

The diazide product (1 eq.) was dissolved in a mixture of EtOAc/THF/1N aq. HCl (5:1:5). Triphenylphosphine (1 eq.) dissolved in diethyl ether was added slowly and the reaction was stirred at room temperature overnight. The white solid was filtered off. 4 N aq. HCl was added to the filtrate and the aqueous phase was washed with diethyl ether, basified with NaOH and subsequently extracted with DCM. The organic phase was dried over Na_2_SO_4_ and the solvent was removed under reduced pressure. The crude product was purified by column chromatography (DCM/MeOH+1%NH_3_ 95:5).

### 2-(2-Azidoethoxy)ethan-1-amine (17a)[65]

The product was synthesized following general procedure C from **16a** (3.36 g, 21.52 mmol, 1 eq.) and triphenylphosphine (5.64 g, 21.52 mmol, 1 eq.). A clear oil was obtained (1.68 g, 60%). R_f_ = 0.38 (DCM/MeOH+1%NH_3_ 95:5).^1^H NMR (300 MHz, CDCl_3_) δ 3.88 (s, 2H), 3.64 – 3.59 (m, 2H), 3.58 – 3.52 (m, 2H), 3.39 – 3.31 (m, 2H), 2.92 (t, *J* = 5.1 Hz, 2H). ^13^C NMR (75 MHz, CDCl_3_) δ 71.47, 69.90, 50.71, 41.06. HRMS (ESI-MS): m/z [M+H]^+^ calculated for C_4_H_11_N_4_O^+^: 131.0927, found 131.0927; C_4_H_10_N_4_O (130.15).

### 2-(2-(2-Azidoethoxy)ethoxy)ethan-1-amine (17b)[50]

The product was synthesized following general procedure C from **16b** (1.93 g, 9.64 mmol, 1 eq.) and triphenylphosphine (2.53 g, 9.64 mmol, 1 eq.). A clear oil was obtained (1.16 g, 69%). R_f_ = 0.32 (DCM/MeOH+1%NH_3_ 95:5). ^1^H NMR (300 MHz, CDCl_3_) δ 3.70 – 3.57 (m, 6H), 3.53 – 3.46 (m, 2H), 3.41 – 3.33 (m, 2H), 2.91 – 2.78 (m, 2H), 1.55 (s, 2H). ^13^C NMR (75 MHz, CDCl_3_) δ 73.51, 70.68, 70.32, 70.08, 50.68, 41.77. HRMS (ESI-MS): m/z [M+H]^+^ calculated for C_6_H_15_N_4_O_2_^+^: 175.1190, found 175.1188; C_6_H_14_N_4_O_2_ (174.20).

### Chloro-5-(4-((3-(1,3-dioxoisoindolin-2-yl)propyl)sulfonamido)phenyl)-3-methyl-2,3,4,5-tetrahydro-1*H*-benzo[d]azepin-7-yl acetate (18)

*13a* (0.10 g, 0.29 mmol, 1 eq.) and pyridine (70 µl, 0.87 mmol, 3 eq.) were dissolved in chloroform. *14a* (0.13 g, 0.44 mmol, 1.5 eq) was added and the reaction was stirred at 50°C overnight. The solvent was removed under reduced pressure and the crude product was purified by column chromatography (95:5 DCM/MeOH+1%NH_3_). A yellow oil was obtained (0.11 g, 62%). R_f_ = 0.30 (95:5 DCM/MeOH+1%NH_3_). ^1^H NMR (400 MHz, CDCl_3_) δ 7.81 – 7.76 (m, 2H), 7.70 – 7.66 (m, 2H), 7.21 – 7.13 (m, 3H), 7.07 – 7.00 (m, 2H), 6.38 (s, 1H), 4.23 (d, *J* = 8.0 Hz, 1H), 3.84 – 3.72 (m, 2H), 3.26 – 2.68 (m, 7H), 2.48 – 2.33 (m, 4H), 2.29 – 2.13 (m, 5H). ^13^C NMR (101 MHz, CDCl_3_) δ 168.68, 168.26, 145.04, 144.16, 140.21, 135.37, 134.21, 131.83, 130.89, 129.52, 124.15, 123.43, 123.19, 121.27, 62.38, 56.50, 53.48, 49.17, 48.48, 47.46, 36.19, 23.24, 20.65. HRMS (ESI-MS): m/z [M+H]^+^ calculated for C_30_H_31_ClN_3_O_6_S^+^: 596.1617, found 596.1623; C_30_H_30_ClN_3_O_6_S (596.10).

### 3-Amino-*N*-(4-(7-chloro-8-hydroxy-3-methyl-2,3,4,5-tetrahydro-1*H*-benzo[d]azepin-1-yl)phenyl)propane-1-sulfonamide (19)

Hydrazine hydrate (83 µl, 1.70 mmol, 10 eq.) was added to a solution of **18** (0.1 g, 0.17 mmol, 1 eq.) in EtOH and the reaction was heated to reflux overnight. The reaction was cooled to 0°C and the white solid was filtered off. The solvent of the filtrate was evaporated under reduced pressure and the crude product was purified by preparative HPLC. A sticky solid was obtained (25 mg, 23%). ^1^H NMR (400 MHz, MeOD) δ 7.46 – 7.09 (m, 5H), 6.82 – 5.80 (m, 1H), 4.57 (d, *J* = 9.7 Hz, 1H), 3.93 – 3.30 (m, 4H), 3.28 – 3.18 (m, 2H), 3.14 – 2.87 (m, 7H), 2.26 – 2.06 (m, 2H). HRMS (ESI-MS): m/z [M+H]^+^ calculated for C_20_H_27_ClN_3_O_3_S^+^: 424.1456, found 424.1459; C_20_H_26_ClN_3_O_3_S*2 TFA (652.00).

### *E*thyl 5-(*N*-(4-(7-chloro-3-methyl-8-((triisopropylsilyl)oxy)-2,3,4,5-tetrahydro-1*H*-benzo[d]azepin-1-yl)phenyl)sulfamoyl)pentanoate (20)

*13b* (0.50 g, 1.01 mmol, 1 eq.) and pyridine (245 µl, 3.03 mmol, 3 eq.) were dissolved in CHCl_3_. **14b** (0.46 g, 2.02 mmol, 2 eq.) was added and the reaction was stirred at 50°C overnight. DCM was added to the reaction and the organic phase was washed with water and brine, dried over Na_2_SO_4_ and the solvent was removed under reduced pressure. The crude product was purified by flash chromatography (SiO_2_, 0 min to 5 min to 25 min, 98:2 to 98:2 to 9:1, DCM/MeOH). A pale yellow solid was obtained (0.37 g, 56%). R_f_ = 0.20 (DCM/MeOH 95:5). ^1^H NMR (400 MHz, CDCl_3_) δ 7.33 – 7.27 (m, 2H), 7.11 – 7.02 (m, 3H), 6.11 (s, 1H), 4.76 – 4.53 (m, 1H), 4.07 (d, *J* = 7.1 Hz, 2H), 3.63 – 3.30 (m, 3H), 3.16 – 2.95 (m, 3H), 2.86 – 2.73 (m, 1H), 2.73 – 2.52 (m, 4H), 2.34 – 2.23 (m, 2H), 1.91 – 1.62 (m, 4H), 1.20 (t, *J* = 7.1 Hz, 3H), 0.95 – 0.86 (m, 21H). ^13^C NMR (101 MHz, CDCl_3_) δ 172.99, 150.47, 142.51, 137.16, 136.69, 132.18, 130.91, 129.31, 122.63, 120.87, 120.13, 61.86, 60.50, 56.73, 51.24, 46.36, 45.59, 33.62, 23.46, 22.99, 17.76, 14.20, 12.56. HRMS (ESI-MS): m/z [M+H]^+^ calculated for C_33_H_52_ClN_2_O_5_SSi^+^: 651.3049, found 651.3060; C_3_H_51_ClN_2_O_5_SSi (651.38).

### 5-(*N*-(4-(7-Chloro-8-hydroxy-3-methyl-2,3,4,5-tetrahydro-1*H*-benzo[d]azepin-1-yl)phenyl)sulfamoyl)-*N*-(prop-2-yn-1-yl)pentanamide (21)

**20** (0.34 g, 0.52 mmol, 1 eq.) was dissolved in THF and LiOH (37 mg, 1.56 mmol, 3 eq.) dissolved in water was added and the reaction was stirred at room temperature overnight. 0.1 N HCl (32 ml, 3.12 mmol, 6 eq.) was added slowly at 0°C. The solvent were removed by lyophilization and the residue was dissolved in DMF. HATU (236 mg, 0.62 mmol, 1.2 eq.), DIPEA (269 µl, 1.56 mmol, 6 eq.) and propargylamine (50 µl, 0.78 mmol, 1.5 eq.) were added and the reaction was stirred at room temperature overnight. The solvent was removed under reduced pressure and the crude product was purified by flash chromatography (SiO_2_, 0 min to 5 min to 6 min to 25 min, 100:0 to 100:0 to 95:5 to 8:2, DCM/MeOH). A yellow solid was obtained (0.24 g, 92%). R_f_ = 0.18 (DCM/MeOH 9:1). ^1^H NMR (400 MHz, MeOD) δ 7.30 – 7.23 (m, 2H), 7.19 – 7.10 (m, 3H), 6.23 (s, 1H), 4.51 (d, *J* = 9.6 Hz, 1H), 3.87 (d, *J* = 2.5 Hz, 2H), 3.63 – 3.36 (m, 3H), 3.25 – 3.16 (m, 1H), 3.14 – 3.05 (m, 2H), 2.97 – 2.85 (m, 2H), 2.82 – 2.74 (m, 3H), 2.53 (t, *J* = 2.6 Hz, 1H), 2.24 – 2.13 (m, 2H), 1.83 – 1.71 (m, 2H), 1.71 – 1.57 (m, 2H). ^13^C NMR (101 MHz, MeOD) δ 173.62, 151.81, 141.96, 137.39, 130.66, 130.42, 129.14, 120.47, 118.31, 116.68, 79.22, 70.86, 60.74, 56.28, 50.55, 45.41, 44.97, 34.63, 31.15, 28.04, 23.84, 22.80. HRMS (ESI-MS): m/z [M+H]^+^ calculated for C_25_H_31_ClN_3_O_4_S^+^: 504.1718, found 504.1722; C_25_H_30_ClN_3_O_4_S (504.04).

### General procedure D

The alkyne (1 eq.) and the linker (1.5 eq.) were dissolved in DCM/MeOH (4:1). CuSO_4_*5 H_2_O (0.1 eq.) and ascorbic acid (0.3 eq.) were added and the reaction was stirred at room temperature for 72 h. The solvents were removed under reduced pressure and the crude product was purified by HPLC.

### 5-(*N*-(4-(7-Chloro-8-hydroxy-3-methyl-2,3,4,5-tetrahydro-1*H*-benzo[d]azepin-1-yl)phenyl)sulfamoyl)-*N*-((1-(2-(2-hydroxyethoxy)ethyl)-1*H*-1,2,3-triazol-4-yl)methyl)pentanamide (22a)

The product was synthesized following general procedure D from **21** (25 mg, 49.60 µmol, 1 eq.), **17a** (9.7 mg, 74.40 µmol, 1.5 eq.), CuSO_4_*5 H_2_O (1.2 mg, 4.96 µmol, 0.1 eq.) and ascorbic acid (2.62 mg, 14.88 µmol, 0.3 eq.). A white solid was obtained (7 mg, 14%). RP-HPLC: >97%, (t_R_ = 7.34 min, k = 1.29). ^1^H NMR (400 MHz, MeOD) δ 7.89 (s, 1H), 7.41 – 7.09 (m, 5H), 6.16 (s, 1H), 4.65 – 4.48 (m, 3H), 4.39 (s, 2H), 3.95 – 3.32 (m, 8H), 3.18 – 2.99 (m, 6H), 2.96 (s, 3H), 2.24 (t, *J* = 6.9 Hz, 2H), 1.87 – 1.61 (m, 4H). HRMS (ESI-MS): m/z [M+H]^+^ calculated for C_29_H_41_ClN_7_O_5_S^+^: 634.2573, found 634.2578; C_29_H_40_ClN_7_O_5_S* 3 TFA (976.26).

### *N*-((1-(2-(2-(2-Aminoethoxy)ethoxy)ethyl)-1*H*-1,2,3-triazol-4-yl)methyl)-5-(*N*-(4-(7-chloro-8-hydroxy-3-methyl-2,3,4,5-tetrahydro-1*H*-benzo[d]azepin-1-yl)phenyl)sulfamoyl)pentanamide (22b)

The product was synthesized following general procedure D from **21** (25 mg, 49.60 µmol, 1 eq.), **17b** (13 mg, 74.40 µmol, 1.5 eq.), CuSO_4_*5 H_2_O (1.2 mg, 4.96 µmol, 0.1 eq.) and ascorbic acid (2.62 mg, 14.88 µmol, 0.3 eq.). A white solid was obtained (4 mg, 8%). RP-HPLC: >97%, (t_R_ = 7.50 min, k = 1.34). ^1^H NMR (400 MHz, MeOD) δ 7.87 (s, 1H), 7.42 – 7.08 (m, 5H), 6.16 (s, 1H), 4.63 – 4.46 (m, 3H), 4.40 (s, 2H), 3.95 – 3.44 (m, 12H), 3.19 – 2.98 (m, 6H), 2.96 (s, 3H), 2.24 (t, *J* = 6.9 Hz, 2H), 1.85 – 1.66 (m, 4H). HRMS (ESI-MS): m/z [M+H]^+^ calculated for C_31_H_45_ClN_7_O_6_S^+^: 678.2835, found 678.2841; C_31_H_43_ClN_6_O_7_S* 3 TFA (1020.32).

### General procedure *E*

The primary amine (1.5 eq) was dissolved in DMF (30 μL). NEt_3_ (10 eq.) and the fluorescent dye NHS-ester (1 eq.) in DMF (60 μL) were added, and the reaction was shaken for 2.5 h in the dark at room temperature. The reaction was quenched with 10% aqueous TFA (20 μL), and the crude product was purified by preparative HPLC.

### 5-((3-(*N*-(4-(7-Chloro-8-hydroxy-3-methyl-2,3,4,5-tetrahydro-1*H*-benzo[d]azepin-1-yl)phenyl)sulfamoyl)propyl)carbamoyl)-2-(6-(dimethylamino)-3-(dimethyliminio)-3*H*-xanthen-9-yl)benzoate (23)

The product was synthesized following general procedure E from **19** (1.86 mg, 2.85 μmol, 1.5 eq), NEt_3_ (2.53 µl, 19 μmol, 10 eq.) and 5-carboxytetramethylrhodamine succinimidyl ester (5-TAMRA NHS ester) (1.00 mg, 1.90 μmol, 1 eq.). A pink solid was obtained (1.6 mg, 89%). RP-HPLC: >97%, (t_R_ = 11.27 min, k = 2.51). HRMS (ESI-MS): m/z [M+H]^+^ calculated for C_45_H_47_ClN_5_O_7_S^+^: 836.2879, found 836.2884; C_45_H_46_ClN_5_O_7_S* TFA (950.28).

### (*E*)-1-(6-((3-(*N*-(4-(7-Chloro-8-hydroxy-3-methyl-2,3,4,5-tetrahydro-1*H*-benzo[d]azepin-1-yl)phenyl)sulfamoyl)propyl)amino)-6-oxohexyl)-2-((*E*)-3-(1-(2-methoxyethyl)-3-methyl-5-sulfo-3-(3-sulfopropyl)-3*H*-indol-1-ium-2-yl)allylidene)-3-methyl-3-(3-sulfopropyl)indoline-5-sulfonate (24)

The product was synthesized following general procedure E from **19** (0.25 mg, 0.38 µmol, 2 eq.) NEt_3_ (0.25 µl, 1.90 µmol, 10 eq.), DY-549P1-NHS-ester (0.2 mg, 0.19 µmol, 1 eq.). A pink solid was obtained (0.1 mg, 39%). RP-HPLC: >97%, (t_R_ 3.59 min, k = 0.21). HRMS (ESI-MS): m/z [M-2H]^2-^ calculated for C_56_H_70_ClN_5_O_17_S^2-^: 639.6535, found 639.6545; C_56_H_72_ClN_5_O_17_S*3 NH_3_ (1335.06).

### 2-(3,6-Bis(dimethylamino)xanthylium-9-yl)-5-((2-(2-(4-((5-(*N*-(4-(7-chloro-8-hydroxy-3-methyl-2,3,4,5-tetrahydro-1*H*-benzo[d]azepin-1-yl)phenyl)sulfamoyl)pentanamido)methyl)-1*H*-1,2,3-triazol-1-yl)ethoxy)ethyl)carbamoyl)benzoate (25)

The product was synthesized following general procedure E from **22a** (3.16 mg, 3.24 μmol, 1.5 eq), NEt_3_ (3.04 µl, 21.60 μmol, 10 eq.) and 5-carboxytetramethylrhodamine succinimidyl ester (5-TAMRA NHS ester) (1.14 mg, 2.16 μmol, 1 eq.). A pink solid was obtained (1.4 mg, 51%). RP-HPLC: >98%, (t_R_ = 10.69 min, k = 2.33). HRMS (ESI-MS): m/z [M+H]^+^ calculated for C_56_H_70_ClN_5_O_17_S^+^: 1046.3996, found 1046.3995; C_54_H_60_ClN_9_O_9_S*2 TFA (1274.68).

### 2-((*E*)-3-((*E*)-1-(6-((2-(2-(4-((5-(*N*-(4-(7-Chloro-8-hydroxy-3-methyl-2,3,4,5-tetrahydro-1*H*-benzo[d]azepin-1-yl)phenyl)sulfamoyl)pentanamido)methyl)-1*H*-1,2,3-triazol-1-yl)ethoxy)ethyl)amino)-6-oxohexyl)-3-methyl-5-sulfonato-3-(3-sulfonatopropyl)indolin-2-ylidene)prop-1-en-1-yl)-1-(2-methoxyethyl)-3-methyl-3-(3-sulfonatopropyl)-3*H*-indol-1-ium-5-sulfonate (26)

The product was synthesized following general procedure E from **22a** (0.28 mg, 0.29 µmol, 1.5 eq.), NEt_3_ (0.27 µl, 1.90 µmol, 10 eq.), DY-549P1-NHS-ester (0.2 mg, 0.19 µmol, 1 eq.). A pink solid was obtained (0.14 mg, 47%). RP-HPLC: >96%, (t_R_ = 3.60 min, k = 0.12). HRMS (ESI-MS): m/z [M-2H]^2-^ calculated for C_54_H_61_ClN_9_O_9_S^2-^: 744.7093, found 744.7106; C_65_H_85_ClN_9_O_19_S_5_*3 NH (1544.29).

### 2-(3,6-Bis(dimethylamino)xanthylium-9-yl)-5-((2-(2-(2-(4-((5-(*N*-(4-(7-chloro-8-hydroxy-3-methyl-2,3,4,5-tetrahydro-1*H*-benzo[d]azepin-1-yl)phenyl)sulfamoyl)pentanamido)methyl)-1*H*-1,2,3-triazol-1-yl)ethoxy)ethoxy)ethyl)carbamoyl)benzoate (27)

The product was synthesized following general procedure E from **22b** (2.64 mg, 2.59 μmol, 1.5 eq), NEt_3_ (2.43 µl, 17.30 μmol, 10 eq.) and 5-carboxytetramethylrhodamine succinimidyl ester (5-TAMRA NHS ester) (0.91 mg, 1.73 μmol, 1 eq.). A pink solid was obtained (1.71 mg, 75%). RP-HPLC: >95%, (t_R_ = 10.75 min, k = 2.35). HRMS (ESI-MS): m/z [M+H]^+^ calculated for C_56_H_65_ClN_9_O_10_S^+^: 1090.4258, found 1090.4258; C_56_H_64_ClN_9_O_10_S*2 TFA (1318.74).

### 2-((*E*)-3-((*E*)-1-(6-((2-(2-(2-(4-((5-(*N*-(4-(7-Chloro-8-hydroxy-3-methyl-2,3,4,5-tetrahydro-1*H*-benzo[d]azepin-1-yl)phenyl)sulfamoyl)pentanamido)methyl)-1*H*-1,2,3-triazol-1-yl)ethoxy)ethoxy)ethyl)amino)-6-oxohexyl)-3-methyl-5-sulfonato-3-(3-sulfonatopropyl)indolin-2-ylidene)prop-1-en-1-yl)-1-(2-methoxyethyl)-3-methyl-3-(3-sulfonatopropyl)-3*H*-indol-1-ium-5-sulfonate (28)

The product was synthesized following general procedure E from **22b** (0.30 mg, 0.29 µmol, 1.5 eq.), NEt_3_ (0.27 µl, 1.90 µmol, 10 eq.), DY-549P1-NHS-ester (0.2 mg, 0.19 µmol, 1 eq.). A pink solid was obtained (0.14 mg, 47%). RP-HPLC: >95%, (t_R_ = 3.78 min, k = 0.18). HRMS (ESI-MS): m/z [M+2H]^2+^ calculated for C_67_H_92_ClN_9_O_20_S_5_^2+^: 768.7370, found 768.7386; C_67_H_90_ClN_9_O_20_S_5_*3 NH (1588.34).

### Radioligand competition binding experiments at the dopamine receptors

Cell homogenates containing the D2longR, D3R, and D4.4R were kindly provided by Dr. Lisa Forster, University of Regensburg. Homogenates containing the D1R and D5R were prepared and radioligand binding experiments with cell homogenates were performed as previously described with minor modifications.[66,67] For radioligand competition binding assays homogenates were incubated in BB at a final concentration of 0.3 μg (D1R), 0.3 μg (D2longR), 0.7 μg (D3R), 0.5-1.0 μg (D4.4R), or 0.4 μg (D5R) protein/well. [^3^H]SCH-23390 (D1R (*K*d = 0.23 nM) and D5R (*K*d = 0.20 nM)) was added in final concentrations of 1.0 nM (D1R) and 1.0 nM (D5R). [^3^H]*N*-methylspiperone (D2longR (*K*d = 0.0149 nM), D3R (*K*d = 0.0258 nM), D4.4R (*K*d = 0.078 nM)) was added in final concentrations of 0.05 nM (D2longR, D3R) or 0.1 nM (D4.4R). Non labelled compounds were added in increasing concentrations for the displacement of the radioligands. After incubation of 60 min (D2longR, D3R, and D4.4R) or 120 min (D1R and D5R) at room temperature, bound radioligand was separated from free radioligand through PEI-coated GF/C filters using a Brandel harvester (Brandel Inc., Unterföhring, Germany), filters were transferred to (flexible) 1450-401 96-well sample plates (PerkinElmer, Rodgau, Germany) and after incubation with scintillation cocktail (Rotiszint eco plus, Carl Roth, Karlsruhe, Germany) for at least 3 h, radioactivity was measured using a MicroBeta2 plate counter (PerkinElmer, Waltham, MA, USA). Competition binding curves were fitted using a four-parameter fit (“log(agonist) vs. response-variable slope”). Calculations of p*K_i_* values with SEM and graphical presentations were conducted with GraphPad Prism 9 software (San Diego, CA, USA).

### *G*_s_ heterotrimer dissociation assay

HEK293A cells (Thermo Fisher) were transiently transfected with the G_s_ BRET sensor, G_s_-CASE (https://www.addgene.org/168124/),[51] along with either D_1_R or D_5_R using polyethyleneimine (PEI). Per well of a 96-well plate, 100 μl of freshly resuspended cells were incubated with 100 ng total DNA mixed with 0.3 μl PEI solution (1 mg/ml) in 10 μl Opti-MEM (Thermo Fisher), seeded onto poly-D-lysine (PDL)-pre-coated white, F-bottom 96-well plates (Brand GmbH) and cultivated at 37°C, 5% CO_2_ in penicillin (100 U/ml)/streptomycin (0.1 mg/ml)-, 2 mM L-glutamin- and 10% fetal calf serum (FCS)-supplemented Dulbecco’s modified Eagle’s medium (DMEM; Thermo Fisher). 48 hours after transfection, cells were washed with Hank’s Buffered Salt Solution (HBSS) and incubated with a 1/1000 furimazine (Promega) dilution in HBSS for 2 minutes. Next, the baseline BRET ratio was recorded in three consecutive reads, cells were stimulated with serial dilutions of **25** or vehicle control, and BRET was recorded for another 15 reads. For experiments in antagonist mode, serial dilutions of **25** were added together with furimazine before the experiment and 1 μM dopamine or vehicle control was added after the first three baseline recordings. All experiments were conducted using a ClarioStar Plus Plate reader (BMG Labtech) with a cycle time of 120 seconds, 0.3 seconds integration time and a focal height of 10 mm. Monochromators were used to collect the NanoLuc emission intensity between 430 and 510 nm and cpVenus emission between 500 and 560 nm. BRET ratios were defined as acceptor emission/donor emission. The basal BRET ratio before ligand stimulation (Ratio_basal_) was defined as the average of all three baselaine BRET values. Ligand-induced ΔBRET was calculated for each well as a percent over basal ([(Ratio_stim_ − Ratio_basal_)/Ratio_basal_] × 100). To correct for non-pharmacological effects, the average ΔBRET of vehicle control was subtracted.

### Fluorescence properties

Excitation and emission spectra of **25-28** were recorded in PBS (137 mM NaCl, 2.7 mM KCl, 10 mM Na_2_HPO_4_, 1.8 mM KH_2_PO_4_, pH 7.4) containing 1% BSA (Sigma-Aldrich, Munich, Germany) using a Cary Eclipse spectrofluorometer (Varian Inc., Mulgrave, Victoria, Australia) at 22 °C, using acryl cuvettes (10 x 10 mm, Sarstedt, Nümbrecht, Germany). The slit adjustments (excitation/emission) were 5/10 nm for excitation spectra and 10/5 nm for emission spectra. Net spectra were calculated by subtracting the respective vehicle reference spectrum, and corrected emission spectra were calculated by multiplying the net emission spectra with the respective lamp correction spectrum. The quantum yield of **25-28** was determined according to previously described procedures[52,68] with minor modifications using a Cary Eclipse spectrofluorometer (Varian Inc., Mulgrave, Victoria, Australia) at 22 °C, using acryl cuvettes (10 x 10 mm, Sarstedt, Nümbrecht, Germany) and cresyl violet perchlorate (Biomol GmbH – Life Science Shop, Hamburg, Germany) as a red fluorescent standard. Absorption spectra were recorded with UV/Vis spectroscopy (350-850 nm, scan rate: 300 nm/min, slits: fixed 2 nm) at concentration of 2 µM for cresyl violet (in EtOH, λ_abs,max_ = 575 nm) and **25-28** (in PBS buffer and PBS + 1% BSA, λ_abs,max_ = 559-561 nm). The excitation wavelength for the emission spectra was 527 nm (**16**) or 555 nm (**17**, **20**). The emission wavelength collected during the excitation scan was 630 nm. The quantum yields were calculated for three different slit adjustments (exc./em.): 5/5 nm, 10/5 nm, 10/10 nm. The means of the quantum yields, absorption and emission maxima are presented in Table 3.

### Live cell confocal microscopy at the D_1_R

Confocal images were recorded with kind assistance from Manel Bosch (Universitat de Barcelona). HEK293T cells were transfected based on previously described procedures with the D1R-YFP (C-terminally tagged) with minor modifications.[69,70,22] Cells were seeded in 35 mm wells containing 1.5 cover slips. 48 h after transfection medium was changed to OptiMem media (Gibco) supplemented with 10 mM HEPES. Imaging was performed using a Zeiss LSM880 Laser Scanning Confocal Microscope equipped with a “Plan-Apochromat” 40x/1,3 Oil DIC M27 objective and a photomultiplier tube (PTM) detector. For excitation of **25** an DPSS laser with a wavelength of 561 nm was used. Fluorescence was detected within an emission window of 569-669 nm. Image size was set to 512 x 512 pixels. After adjusting the focus, time-lapse images were recorded in intervals between 0.32 and 1 s. **25** was added in a final concentration of 50 nM. Dissociation was induced by the addition of SCH-23390 (50 µM, 1,000-fold excess). Time-lapse confocal images were processed using the ImageJ software. Contrast was adjusted for each file to facilitate the visualization of the fluorescence signal. Total fluorescence was plotted as a function of time using GraphPad Prism9 software (GraphPad, San Diego, USA). The time of addition of **25** was set as 0 min. Data was fitted using the “Association – one conc. of hot ligand” or “association then dissociation” functions from Prism9. *K_on_*, *K_off_* and kinetic *K_d_* values were calculated by Prism9.

### Molecular Brightness

HEK-293AD cells were seeded in 8-well Ibidi µ-slides with a density of 25,000 cells per well and transfected with 2 μg hD_1_R-mNeonGreen after 24 h using JetPrime transfection reagent according to manufacturer’s protocol. After further 24 h cells were washed and imaged in FRET-buffer (144 mM NaCl, 5.4 mM KCl, 1 mM MgCl_2_, 2 mM CaCl_2_, 10 mM HEPES) on a confocal laser scanning microscope, Leica SP8, with a white-light laser at wavelengths of 488 and 552 nm, and laser power of 5%. All measurements were conducted with an HC PLAP CS2 40 × 1.3 numerical aperture (NA) oil immersion objective (Leica). Movies were acquired at 1.3 seconds per frame for 100 frames with two hybrid detectors in the range of 498 to 547 nm and 562 to 612 nm respectively, in a line sequential, counting mode. Molecular brightness ε and number of molecules N are calculated from the average (k) of the photon counts collected in a pixel and its variance (σ) according to the formulas ε = σ^2^/k-1 and N= k^2^/ σ^2^. ImageJ was used to extract molecular brightness and fluorescence intensity values, number of emitters was calculated with Word Excel and obtained values were plotted and fitted with Prism v. 9.5.1.

## *N*otes

The authors declare no competing financial interest.

## Author contribution

The authors contributed as follows: N.R.: synthesis and analytics of the compounds, radioligand binding studies, fluorescence properties and manuscript writing. D.M.: radioligand binding studies, preparation of mNeonGreen constructs. M.N.: synthesis and radioligand binding studies. H.S.: G_s_ heterotrimer dissociation assay, manuscript writing. A.S.: Molecular Brightness studies, manuscript writing. N.K.: Molecular Brightness studies. I.R.-R.: Cell culture for confocal microscopy. G.N.: Cell culture for confocal microscopy, conception. R.F.: conception, providing infrastructure. P.K.: conception, providing infrastructure. P.A.: conception, manuscript writing, providing infrastructure. S.P.: conception, project administration, manuscript writing, confocal microscopy, providing infrastructure. The data were analyzed and discussed by all authors. All authors have given approval to the final version of the manuscript.

## Supporting Information

The Supporting Information is available free of charge at link DOI: XXX/number

Chemical purity and stability (Figure S1-S6), Dopamine-induced G_s_ activation (Figure S7), Confocal microscopy (Figure S8), NMR spectra (Figure S9-S25), Structures of the fluorescent ligands **23-28** (Figure S26-S31) (PDF).

## Acknowledgements

We thank Manel Bosch (Universitat de Barcelona) for expert technical assistance in confocal microscopy. We thank Dr. Lisa Forster for providing cell homogenates for the D_2long_R, D_3_R and D_4_R. We thank Prof. Dr. Sigurd Elz for providing infrastructure. S.P. was supported by the Fonds der Chemischen Industrie (No. 661688) and University of Regensburg (Academic Research Sabbatical Program). Financial support by the graduate school “Receptor Dynamics: Emerging Paradigms for Novel Drugs (K-BM-2013-247)” of the Elite Network of Bavaria (ENB) for S.P., N.R., M.N. and H.S. is gratefully acknowledged. We would like to thank the Deutsche Forschungsgemeinschaft (DFG) for support through project 421152132 SFB1423 subproject C03 (P.A.) and SFB 1470 subproject A01 (P.A.). R.F., as PI, was funded by Spanish MCIN/AEI/10.13039/501100011033 (grant PID2021-126600OB-I00) and by the European Union Next Generation EU/PRTR (ERDF A way of making Europe).

