## Supporting Information for "Shedding Light on the D_1_-Like Receptors: A Fluorescence-Based Toolbox for Visualization of the D_1_ and D_5_ Receptors"

### Contents


1. Chemical purity and stability ........ S3
2. Dopamine-induced $G_s$ activation ........ S6
3. Confocal microscopy ........ S7
4. NMR spectra ........ S8
5. Structures of the fluorescent ligands **23-28** ........ S17

### 1. Chemical purity and stability

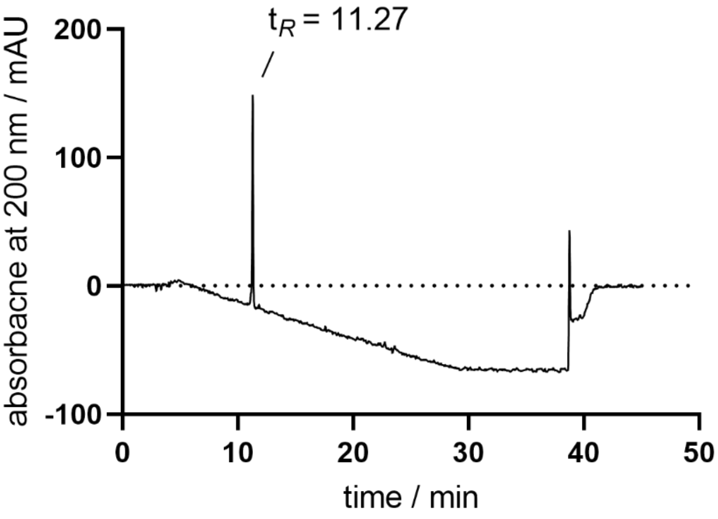


**Figure S1.** RP-HPLC analysis (purity control) of **23** (>97%, 220 nm).

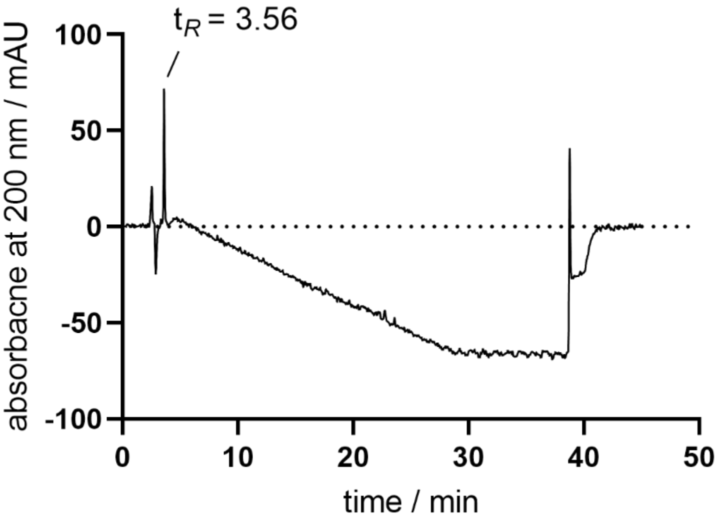


**Figure S2.** RP-HPLC analysis (purity control) of **24** (>97%, 220 nm).

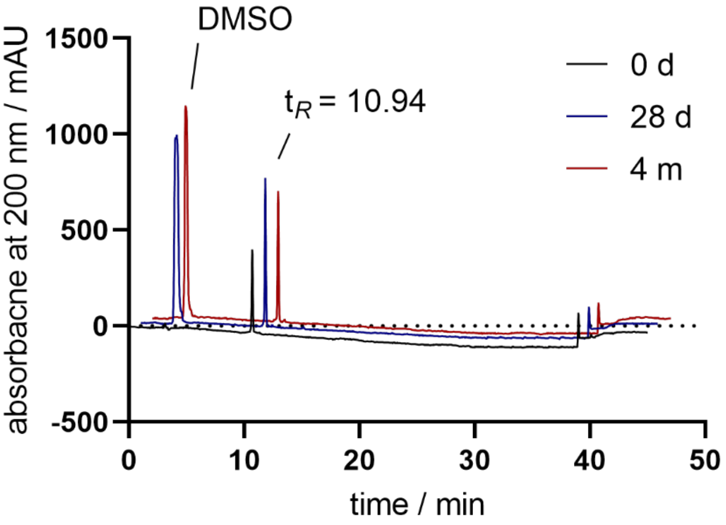


**Figure S3.** RP-HPLC analysis (purity and stability control) of **25** (>98%, 220 nm). Stability at -18°C after 28 days and 4 months.

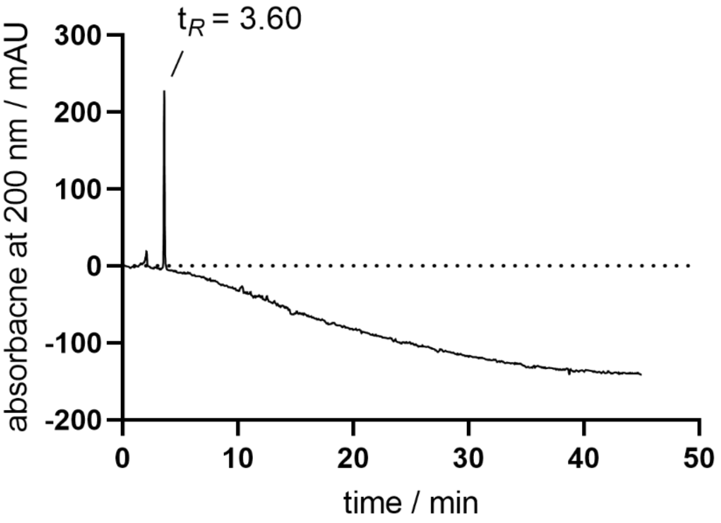


**Figure S4.** RP-HPLC analysis (purity control) of **26** (>96%, 220 nm).

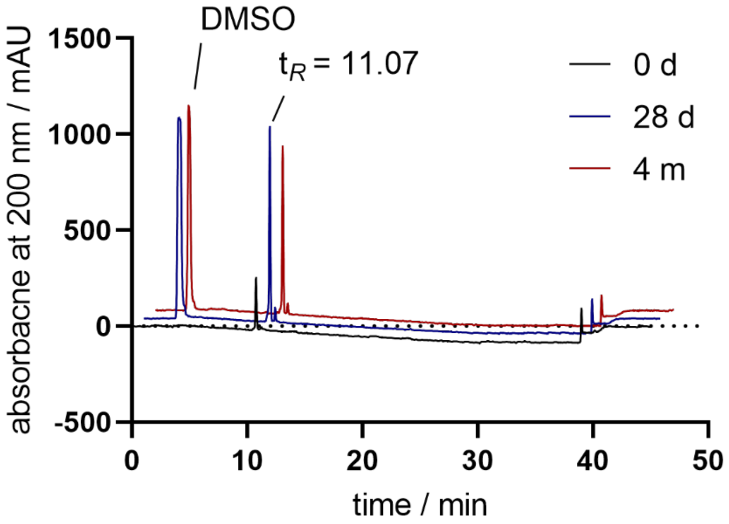


**Figure S5.** RP-HPLC analysis (purity and stability control) of **27** (>95%, 220 nm). Stability at -18°C after 28 days and 4 months.

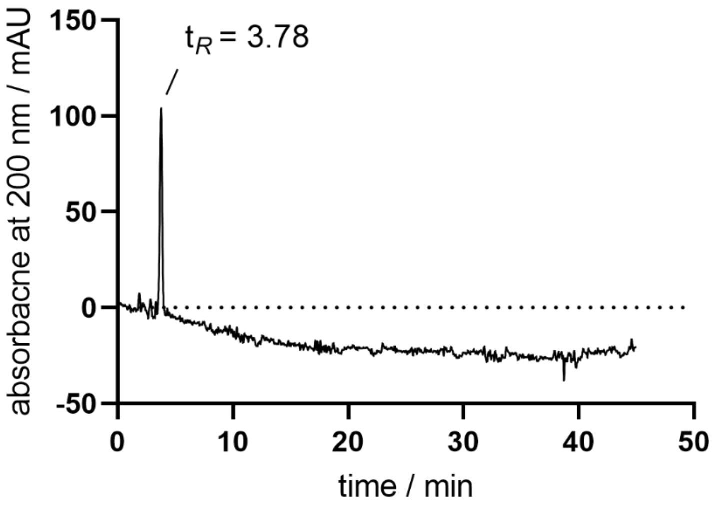


**Figure S6.** RP-HPLC analysis (purity control) of **28** (>95%, 220 nm).

### 2. Dopamine-induced $G_s$ activation

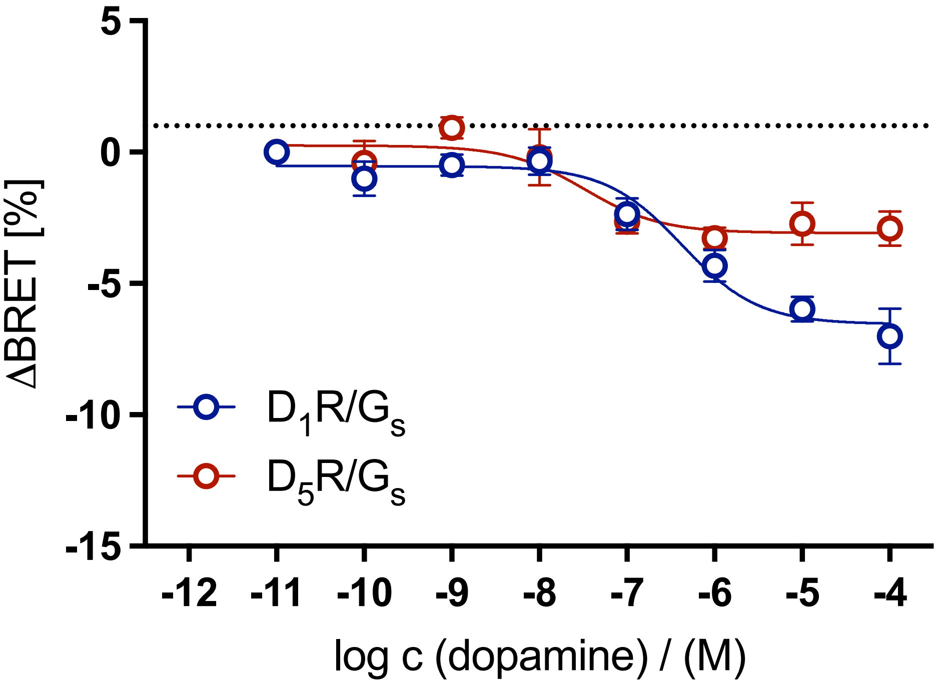


**Figure S7.** Concentration−response curves (CRCs) for $G_s$ activation of dopamine in HEK293A cells transiently expressing the $G_s$ BRET sensor along with the wild-type $D_1R$ or $D_5R$. Graphs represent the means of five ($D_1R$) or four ($D_5R$) independent experiments each performed in duplicate. Data were analyzed by nonlinear regression and were best fitted to sigmoidal concentration-response curves.

### 3. Confocal microscopy

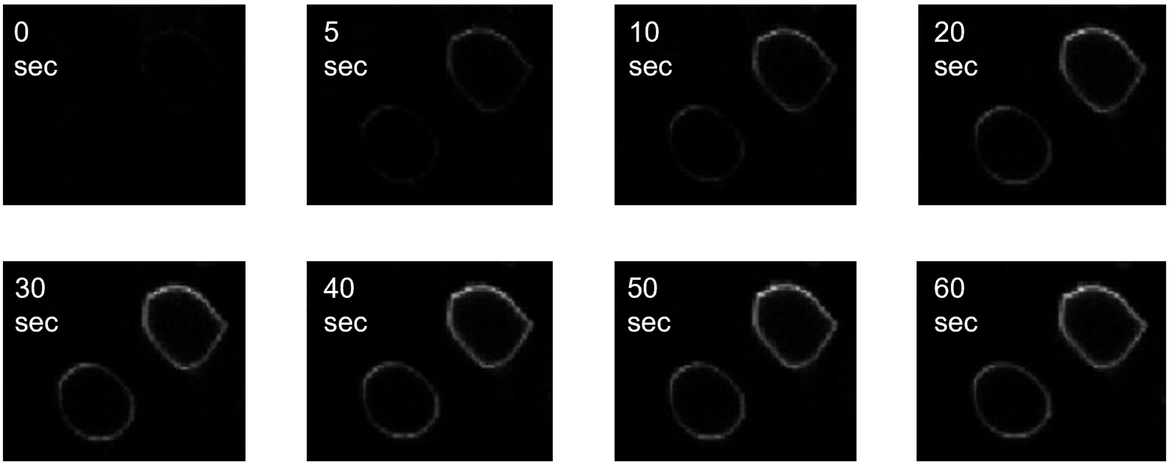


**Figure S8.** Association of **25** to the $hD_1R$ at HEK-293T cells using LSCM. Time-lapse confocal microscopy images of **25** (c = 50 nM) at HEK-293T cells transiently expressing the $hD_1R$ (**A**).

### 4. NMR spectra

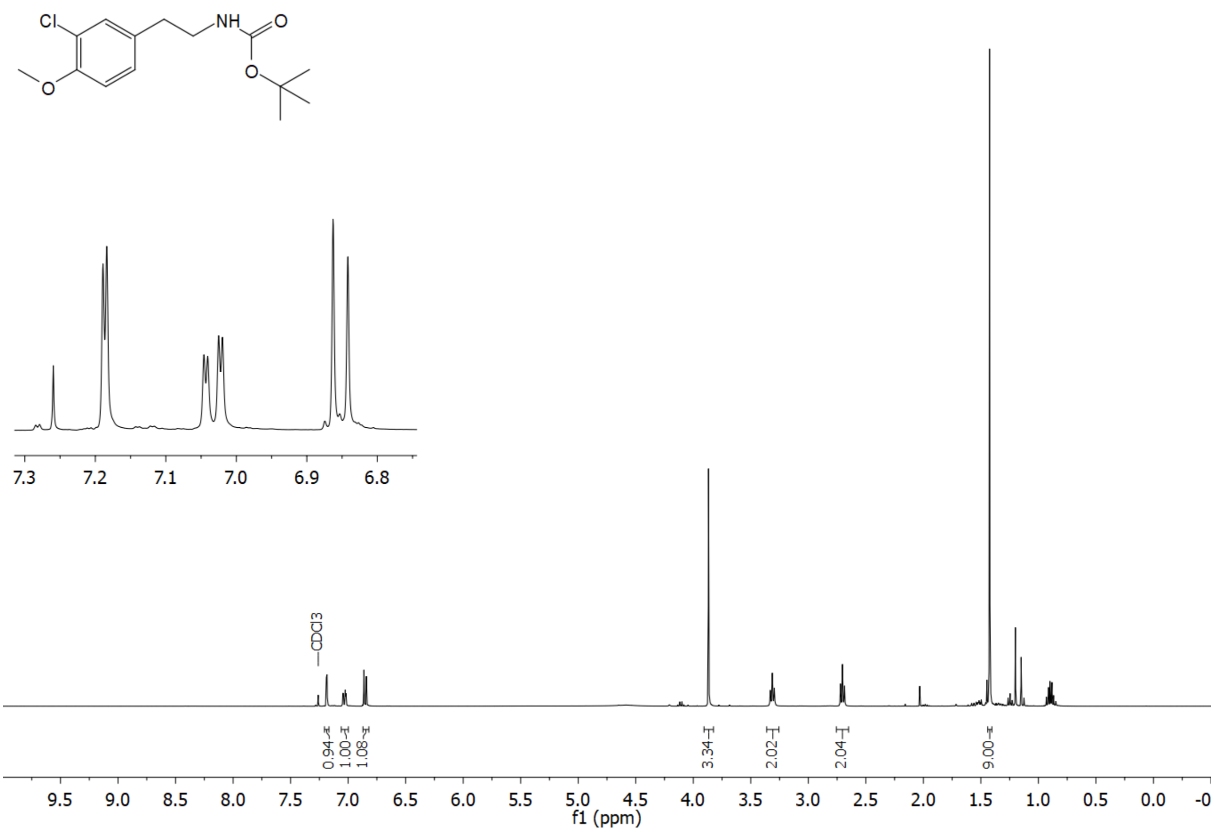


**Figure S9.** $^{1}H$ NMR spectrum (400 MHz, $CDCl_3$) of compound **7**.

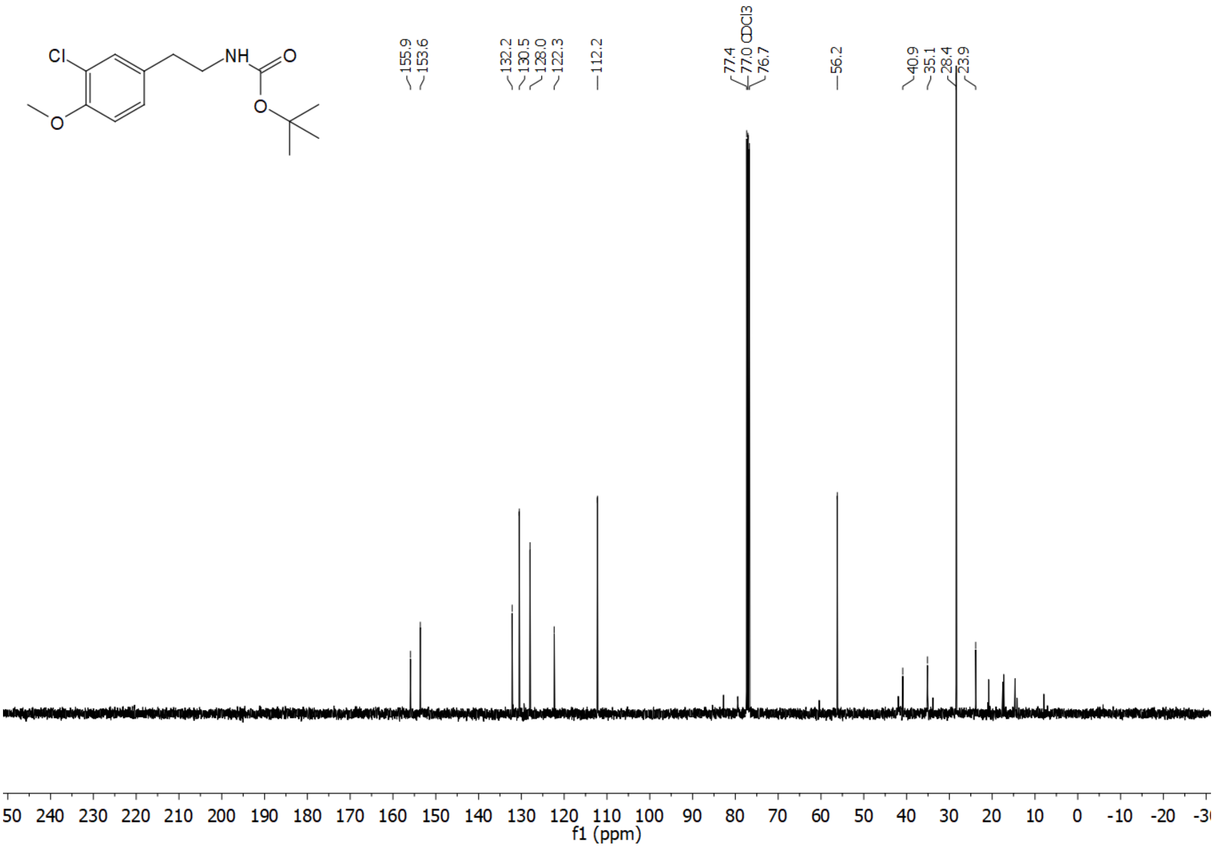


**Figure S10.** $^{13}C$ NMR spectrum (101 MHz, $CDCl_3$) of compound **7**.

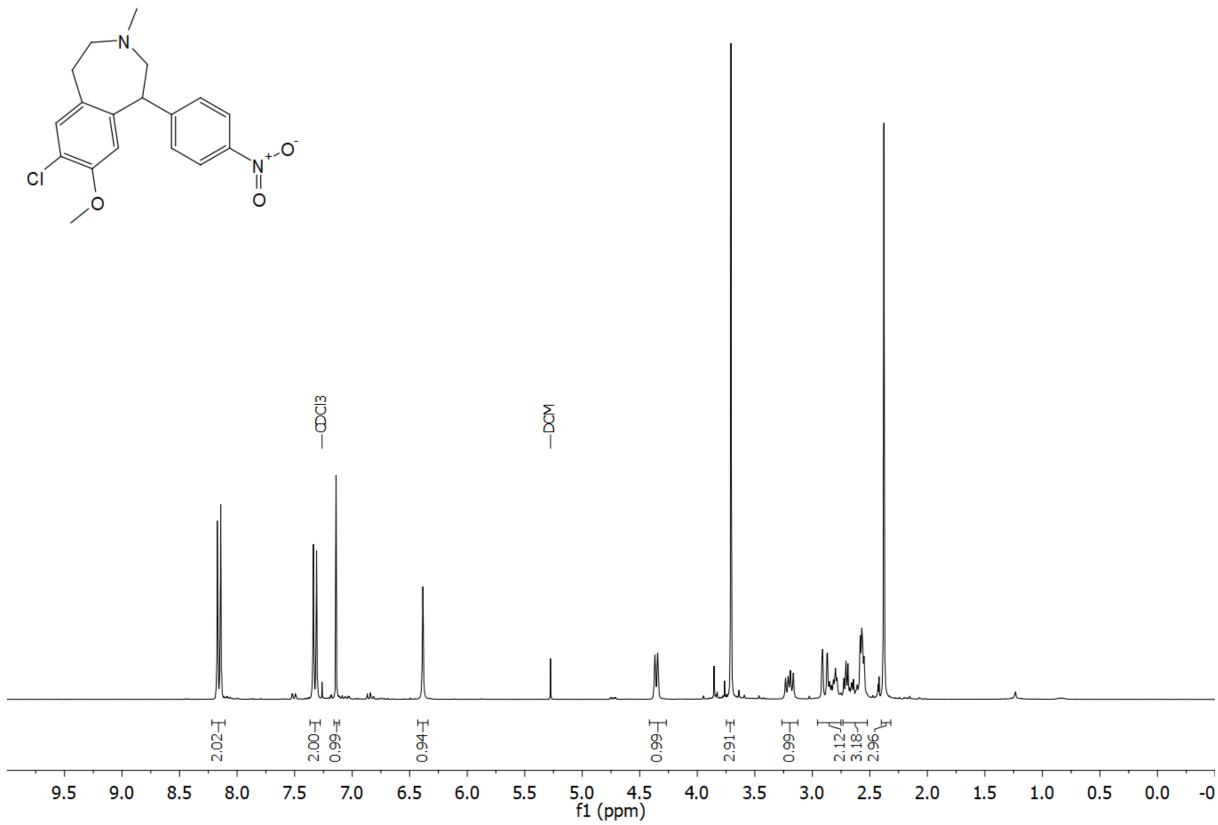


**Figure S11.** $^{1}$H NMR spectrum (300 MHz, $CDCl_3$) of compound **10**.

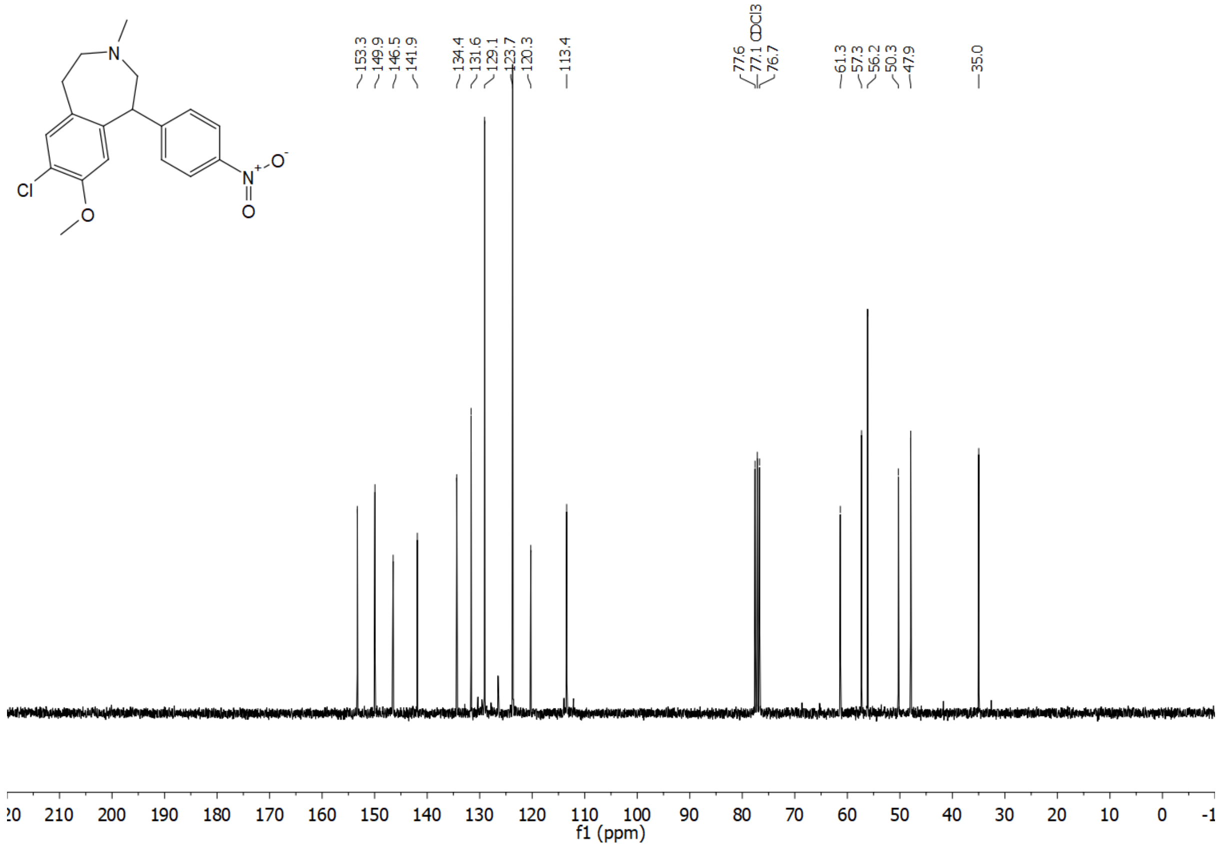


**Figure S12.** $^{13}$C NMR spectrum (75 MHz, $CDCl_3$) of compound **10**.

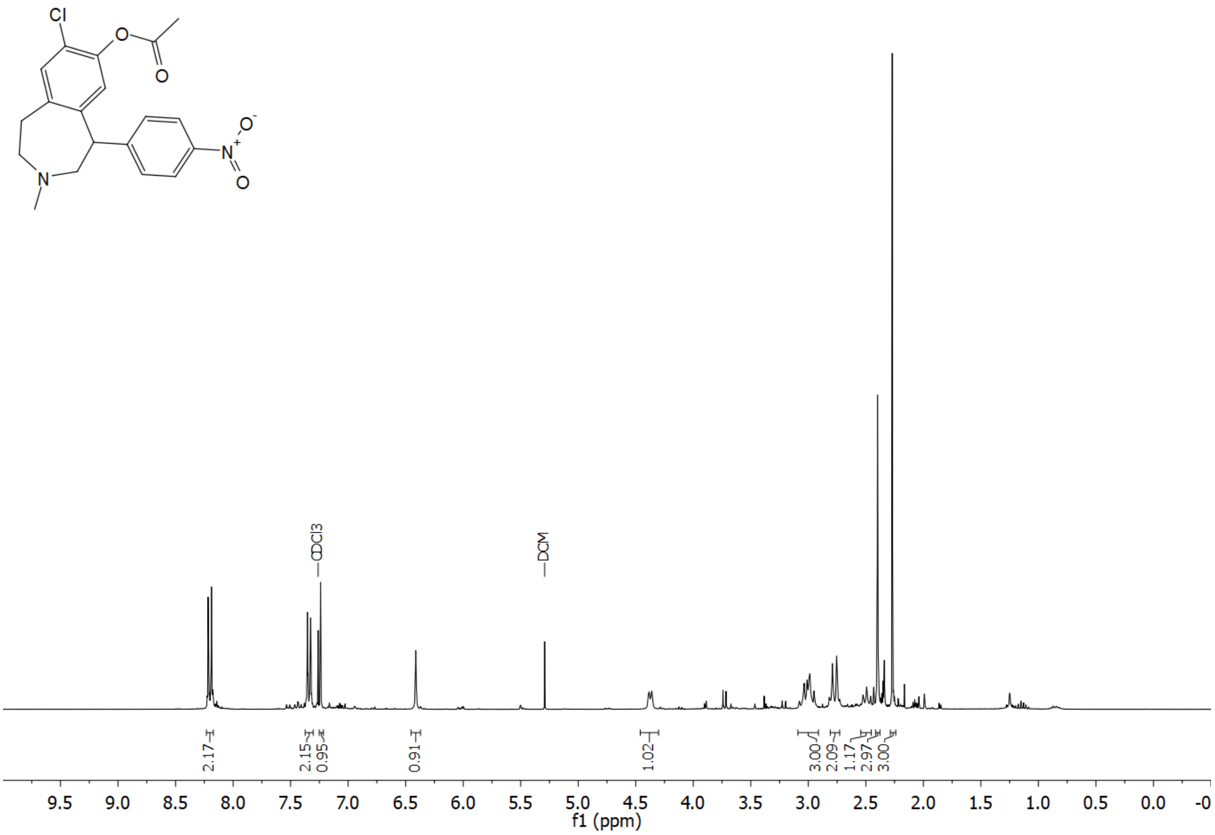


**Figure S13.** $^{1}$H NMR spectrum (300 MHz, $CDCl_3$) of compound **12a**.

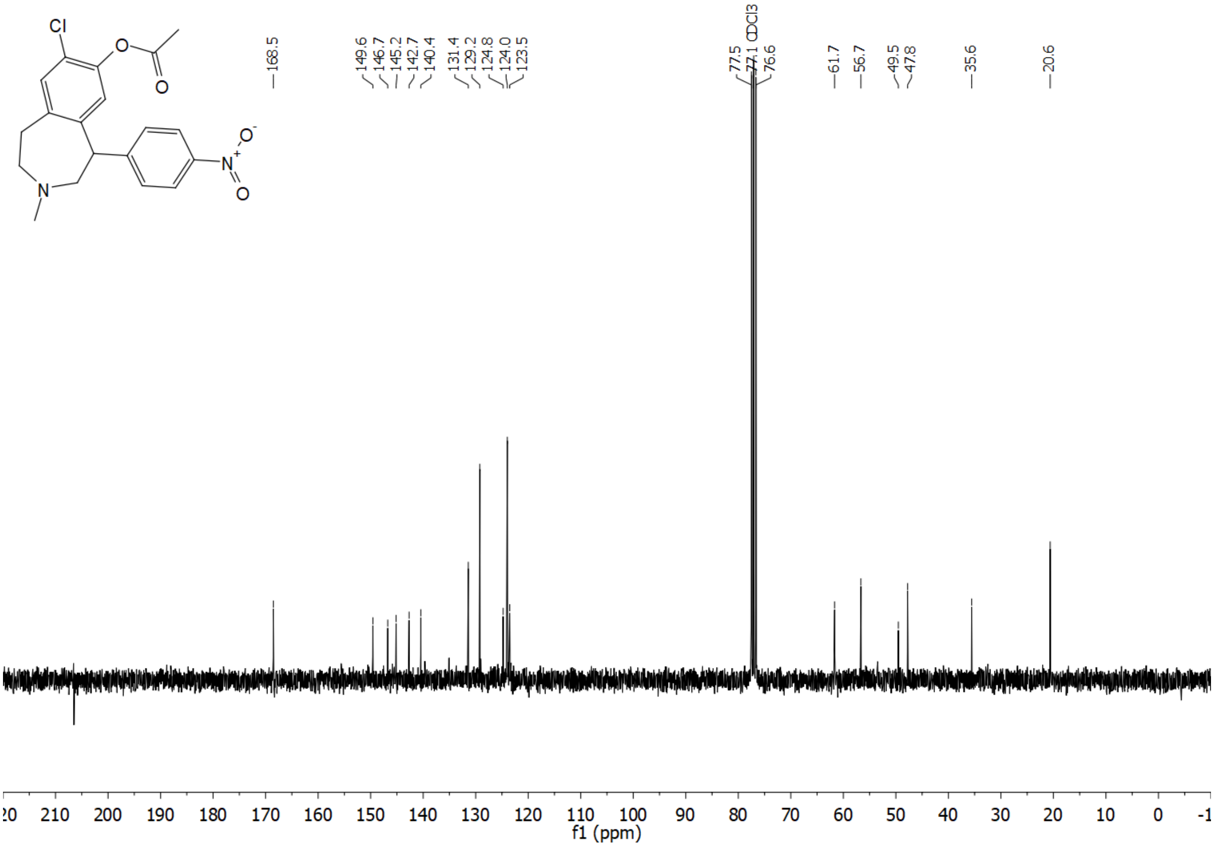


**Figure S14.** $^{13}$C NMR spectrum (75 MHz, $CDCl_3$) of compound **12a**.

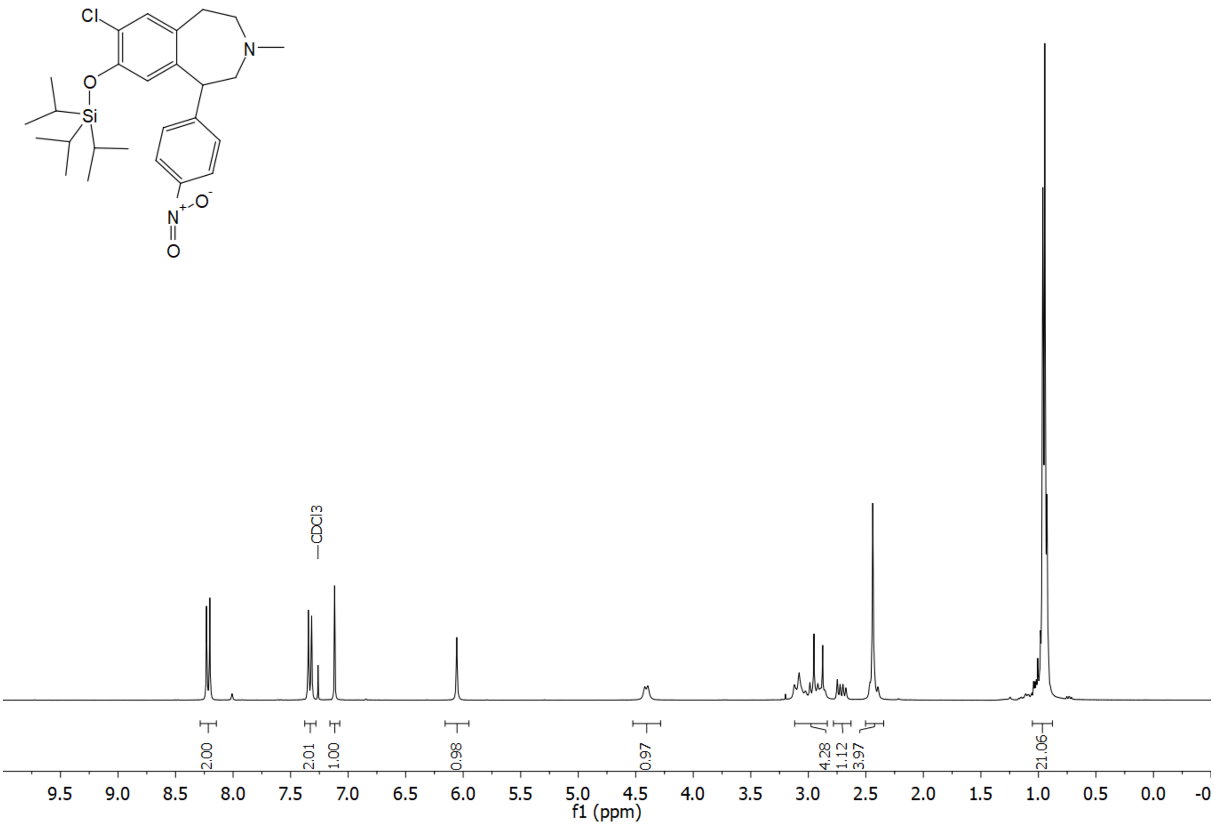


**Figure S15.** $^{1}H$ NMR spectrum (300 MHz, $CDCl_3$) of compound **12b**.

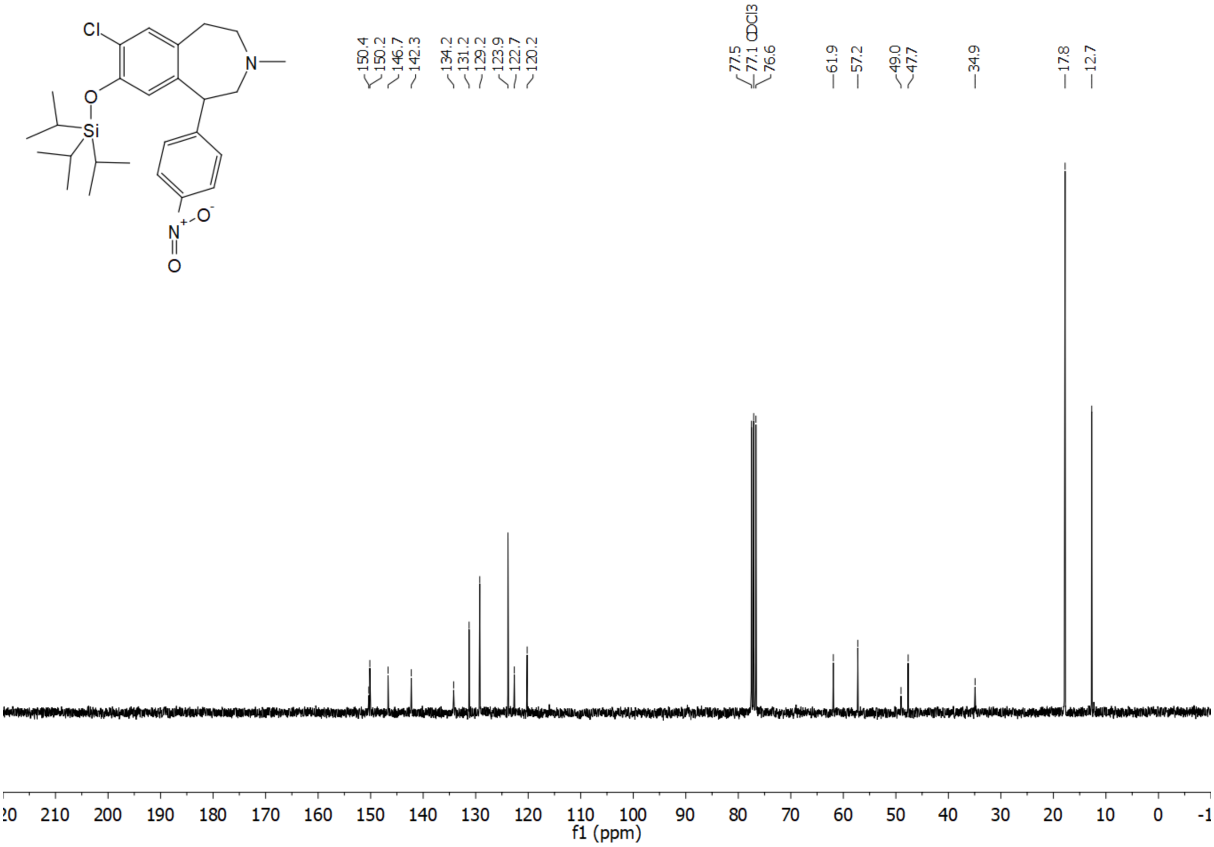


**Figure S16.** $^{13}C$ NMR spectrum (75 MHz, $CDCl_3$) of compound **12b**.

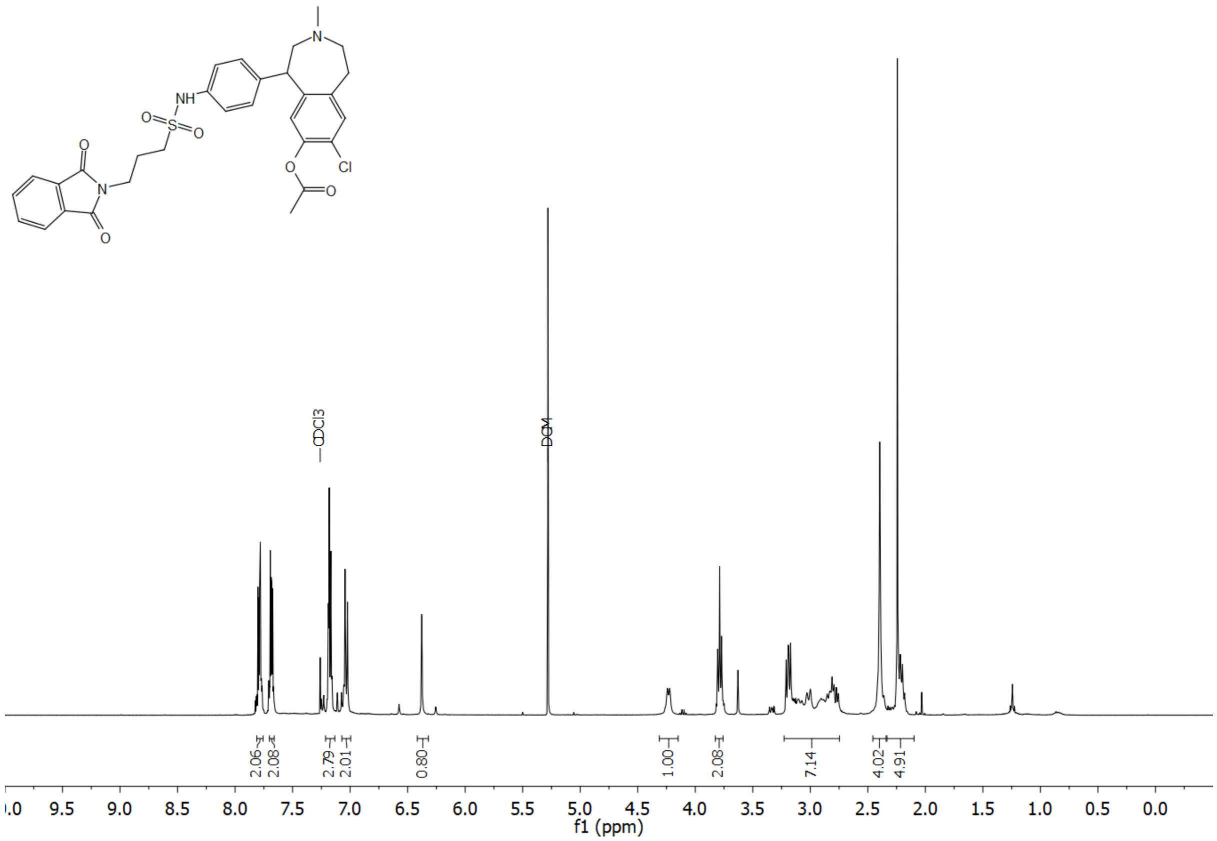

**Figure S17.** $^{1}H$ NMR spectrum (400 MHz, $CDCl_3$) of compound **18**.

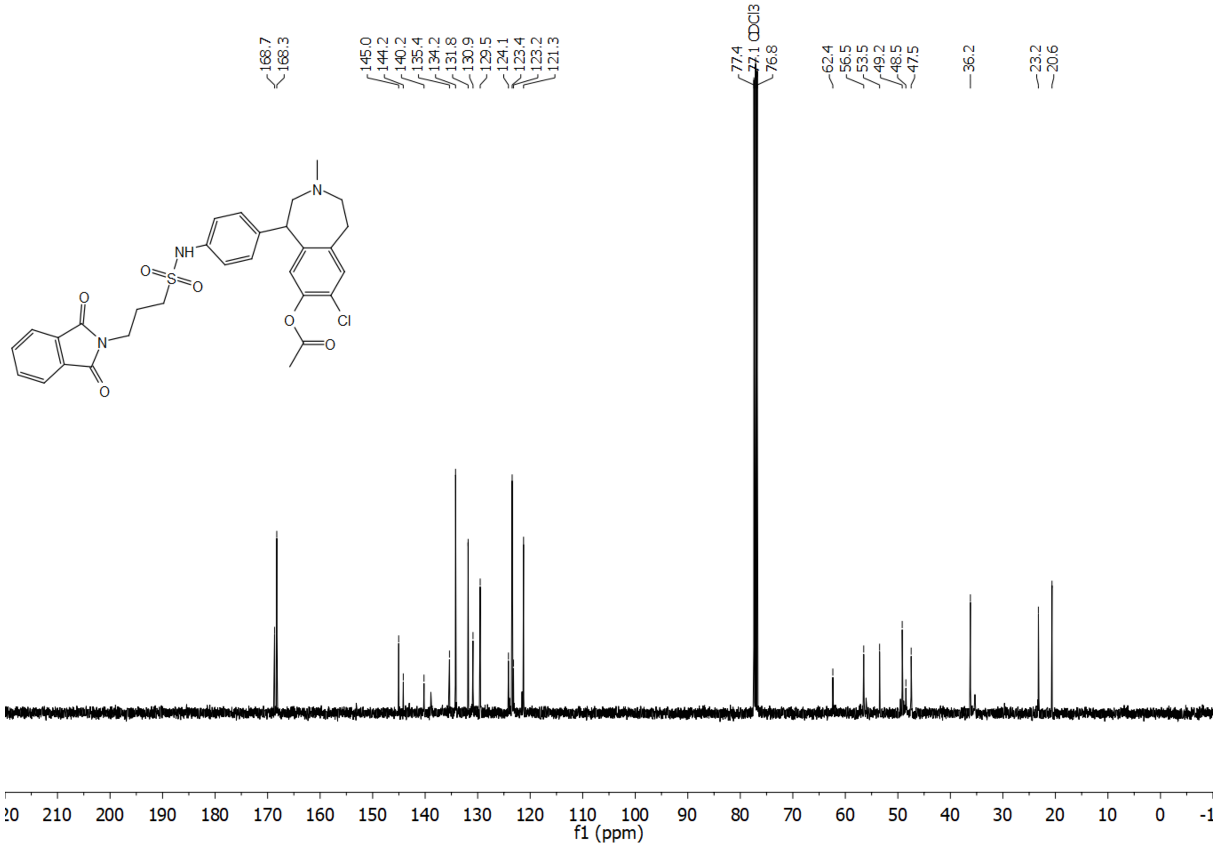

**Figure S18.** $^{13}C$ NMR spectrum (101 MHz, $CDCl_3$) of compound **18**.

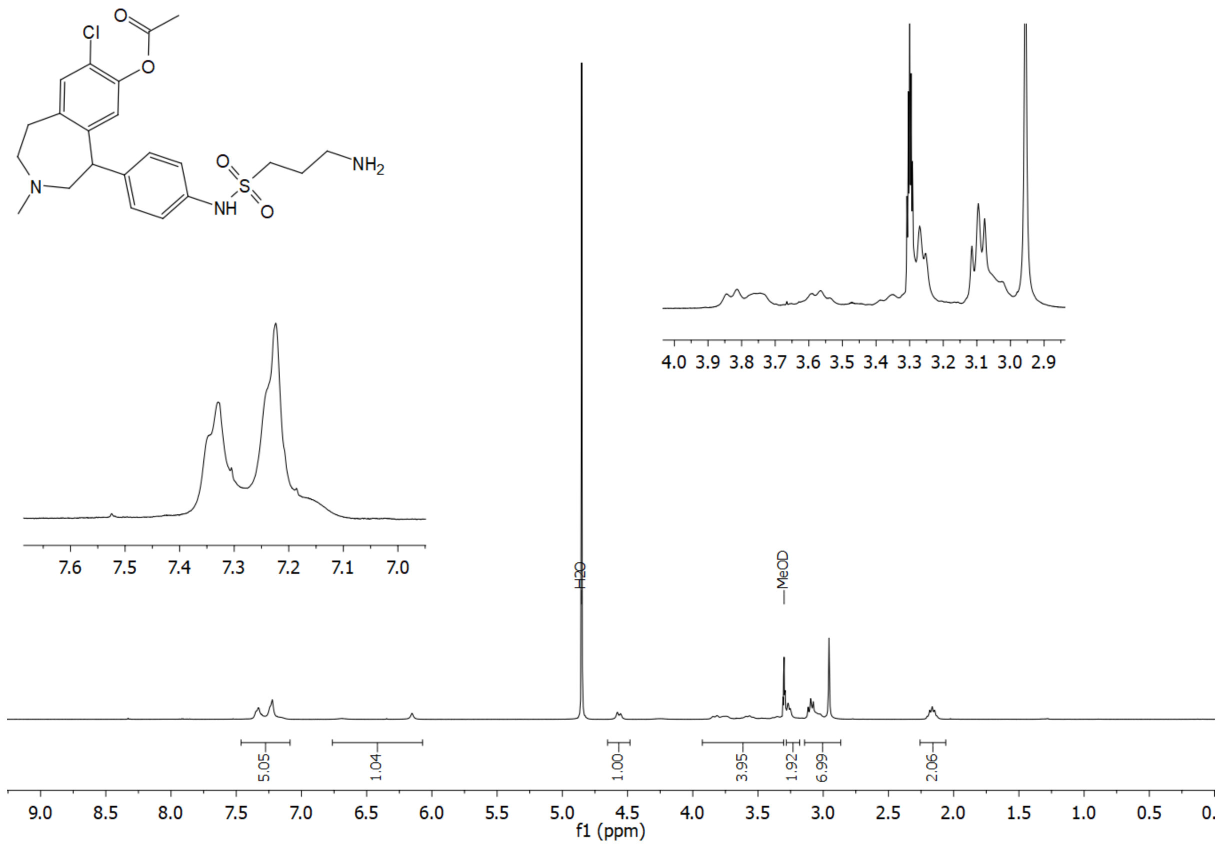


**Figure S19.** $^1$H NMR spectrum (400 MHz, MeOD) of compound **19**.

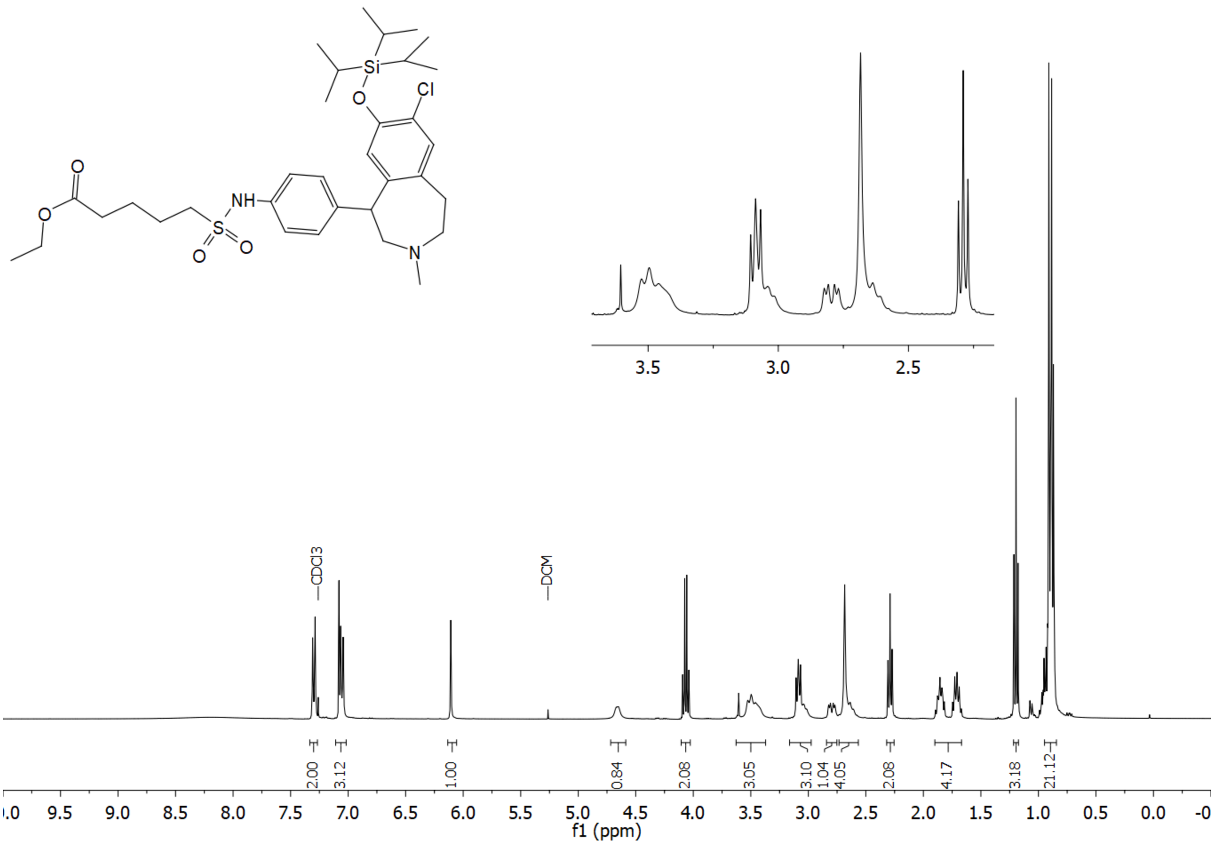


**Figure S20.** $^{1}$H NMR spectrum (400 MHz, $CDCl_3$) of compound **20**.

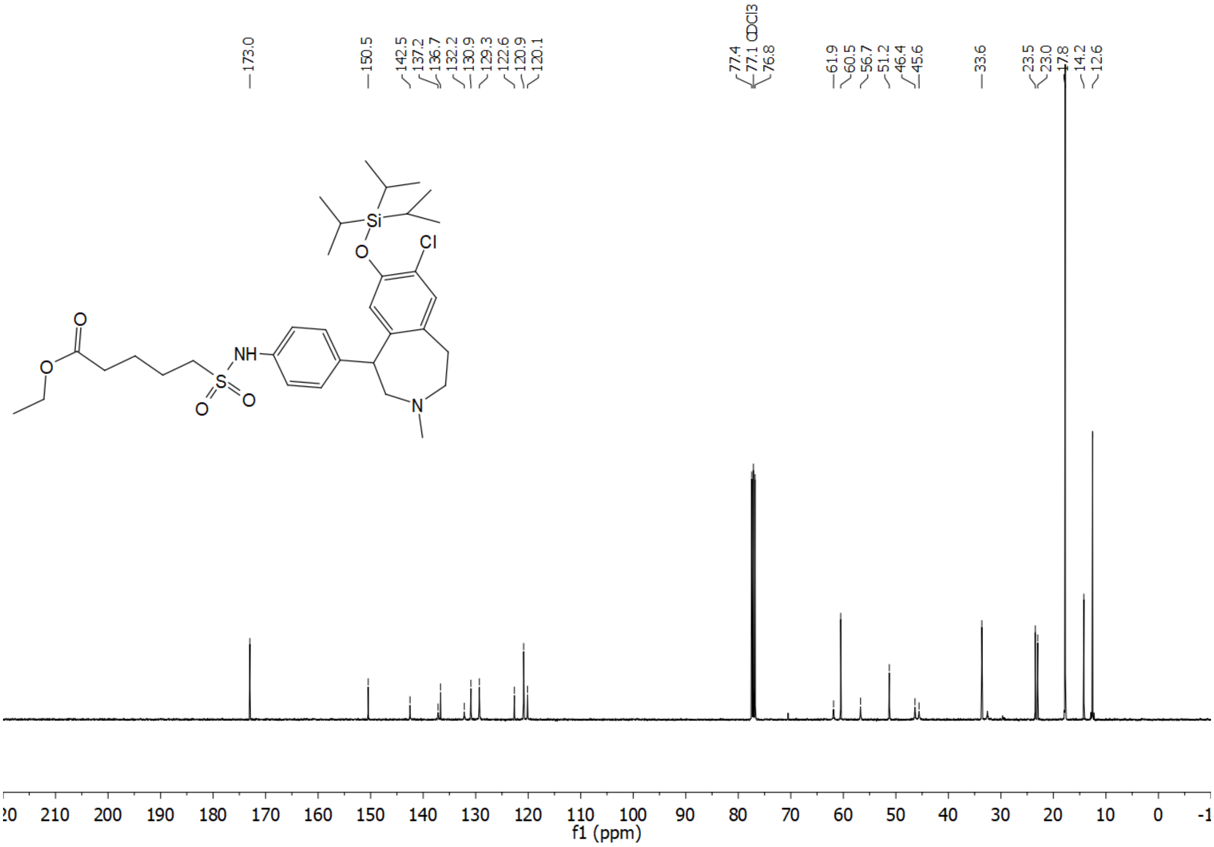


**Figure S21.** $^{13}$C NMR spectrum (101 MHz, $CDCl_3$) of compound **20.**

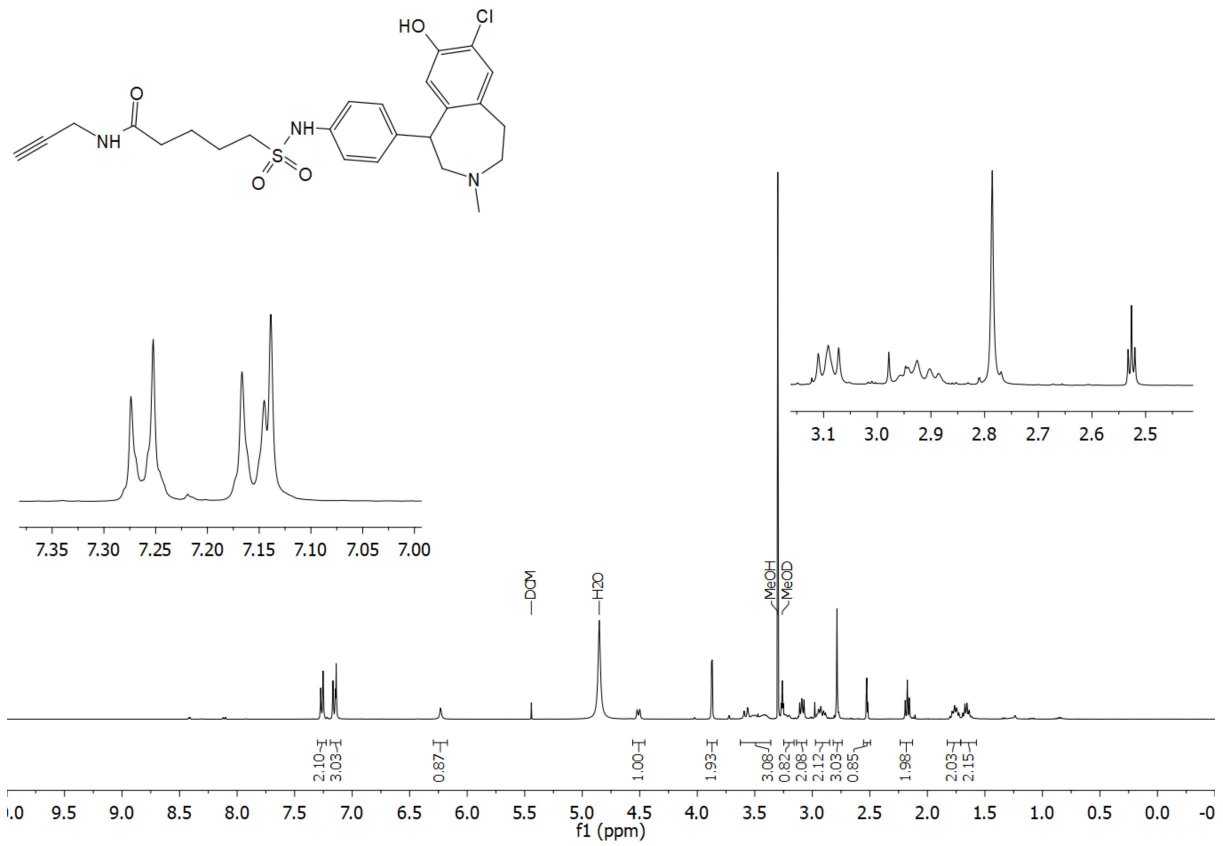

**Figure S22.** $^{1}$H NMR spectrum (400 MHz, MeOD) of compound **21**.

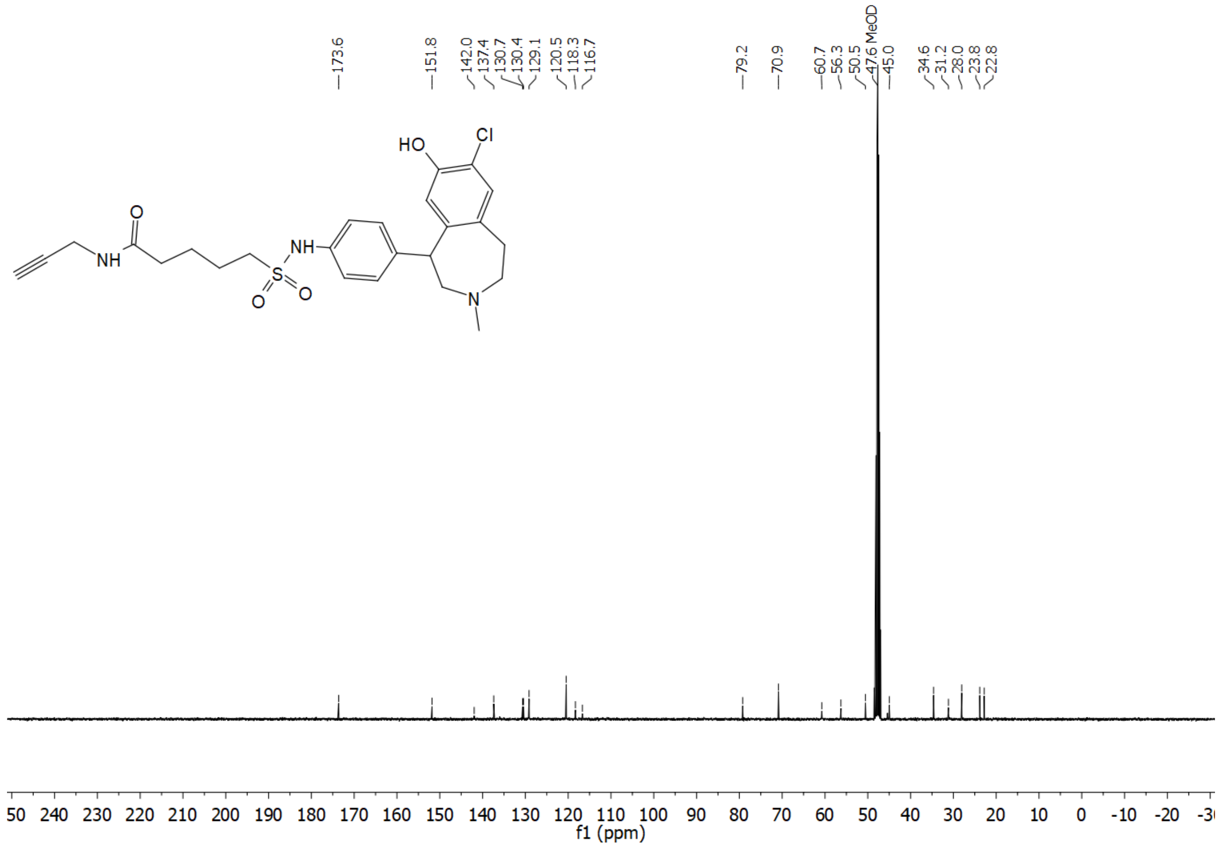

**Figure S23.** $^{13}$C NMR spectrum (101 MHz, MeOD) of compound **21**.

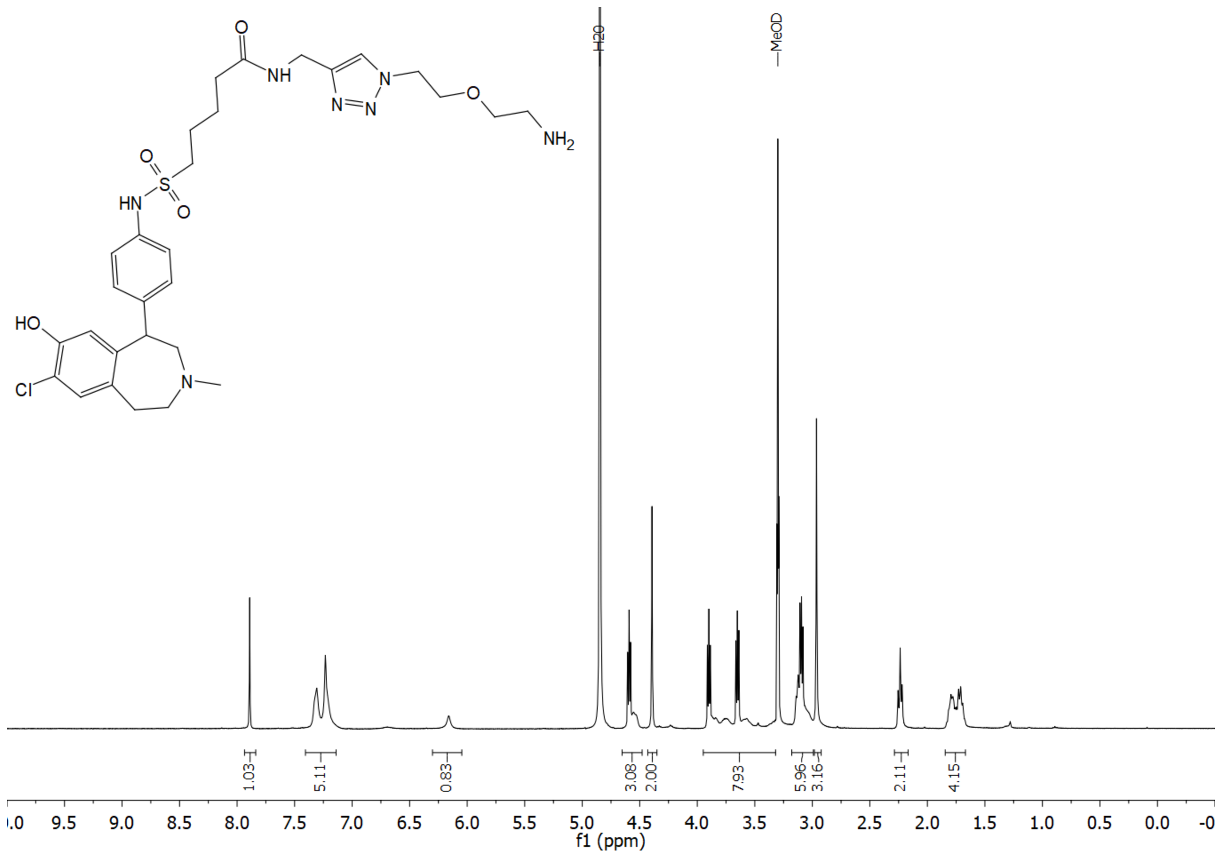

**Figure S24.** $^{1}$H NMR spectrum (400 MHz, MeOD) of compound **22a**.

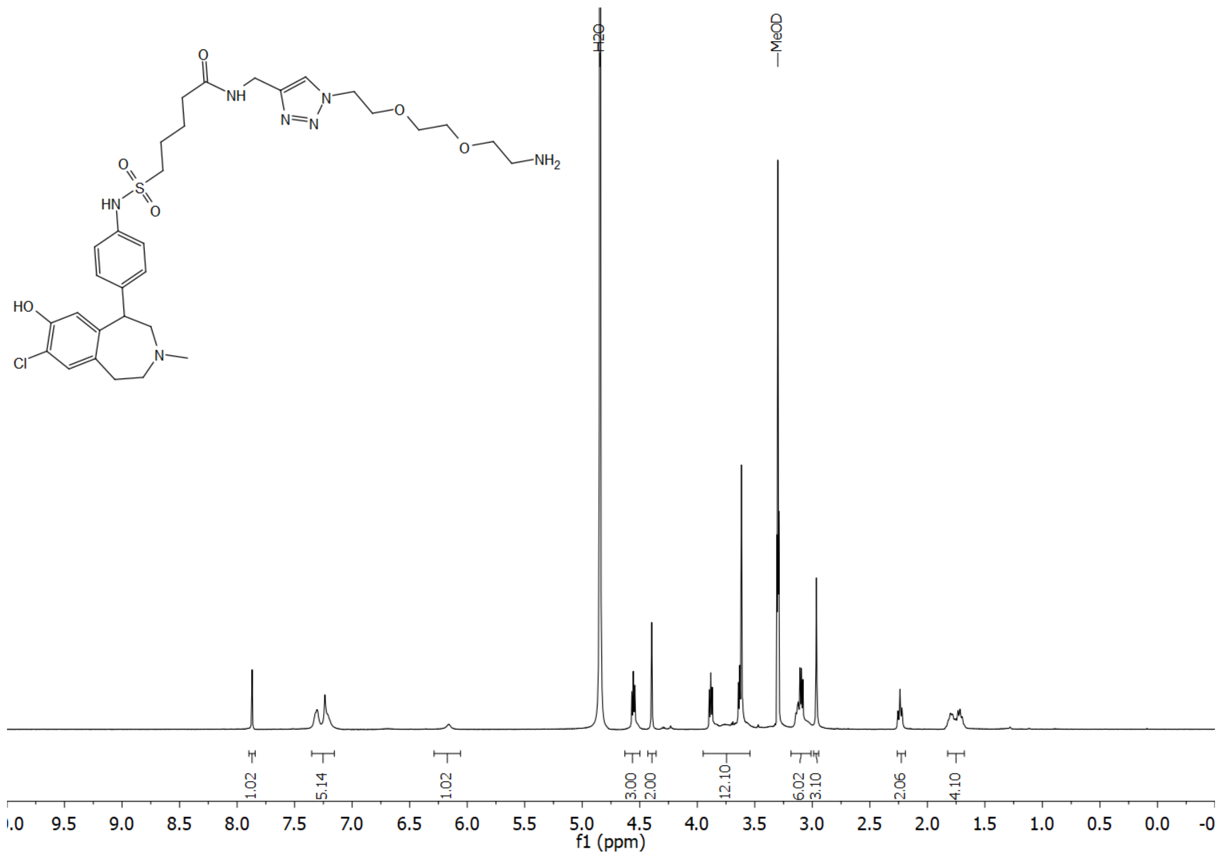

**Figure S25.** $^{1}$H NMR spectrum (400 MHz, MeOD) of compound **22b**.

### 5. Structures of the fluorescent ligands 23-28

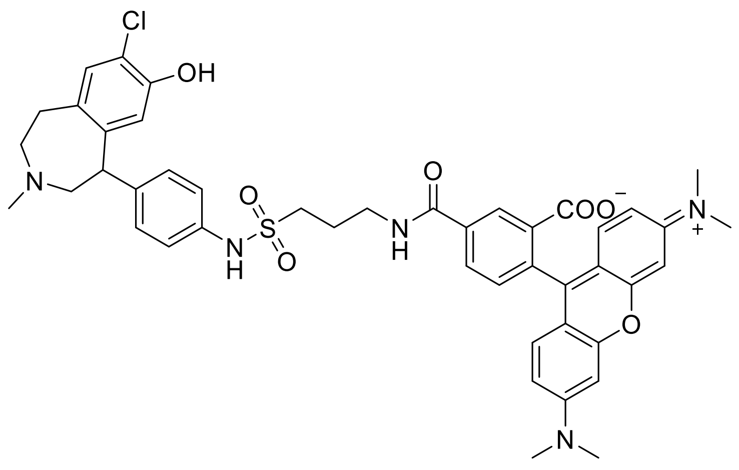


**Figure S26.** Structure of compound **23**.

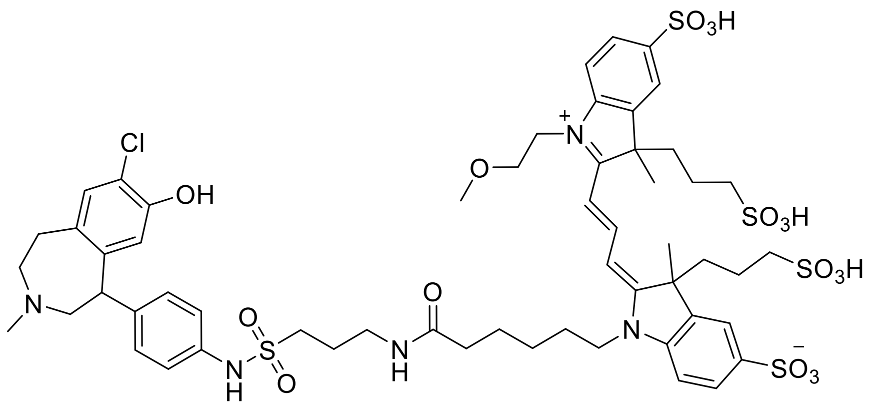


**Figure S27.** Structure of compound **24**.

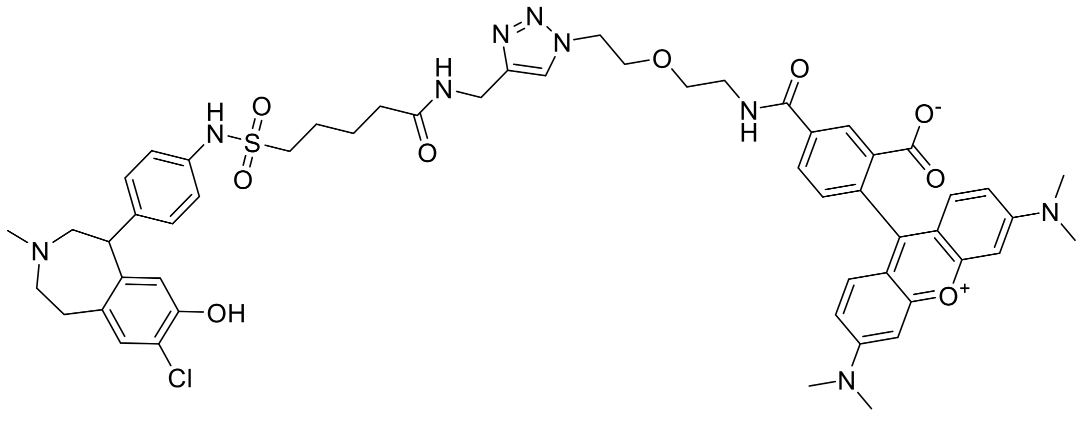


**Figure S28.** Structure of compound **25**.

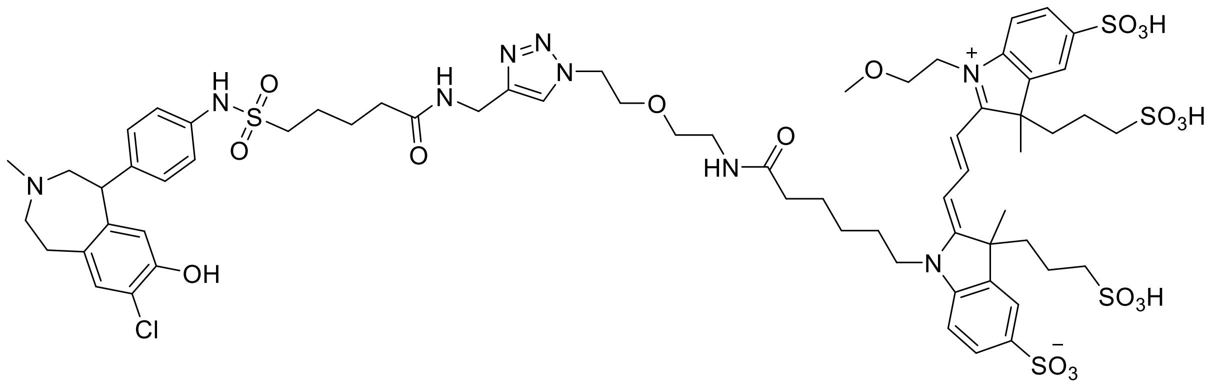


**Figure S29.** Structure of compound **26**.

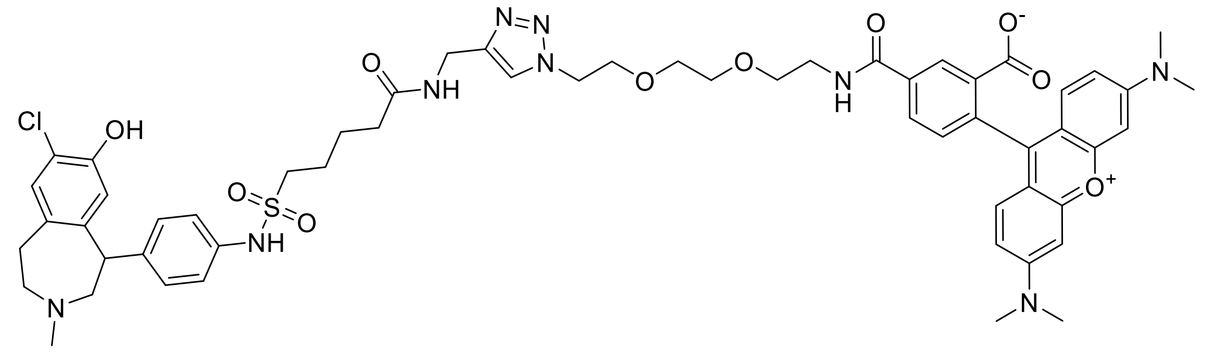


**Figure S30.** Structure of compound **27**.

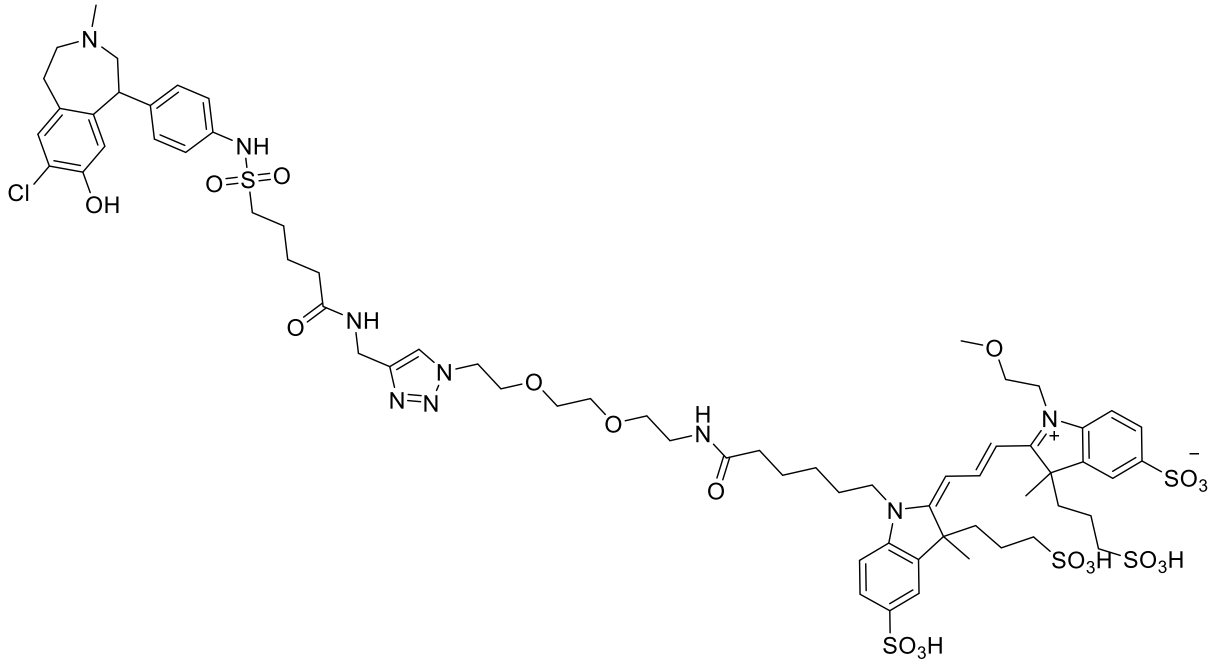


**Figure S31.** Structure of compound **28**.
